# Dincta: Data Integration and Cell Type Annotation of Single Cell Transcriptomes

**DOI:** 10.1101/2020.09.28.316901

**Authors:** Songting Shi

## Abstract

We proposed a method for data integration and cell type annotation (Dincta) of single cell transcriptomes in a unify framework. The Dincta can handle three cases. In the first case, the data has been annotated the cell type for all cells, Dincta can integrate the the data into a common low dimension embedding space such that cells with different cell types separate while cells from the different batches but in the same cell type cluster together. In the second case, the data was only annotated for part of cells, such as one sample, Dincta can integrate the data into a common low dimension embedding space such that cells with different cell types separate while cells from the different batches but in the same cell type cluster together. Moreover, it can infer the known or novel cell type of the cells with unknown cell type initially. In the third case, there are no cell type information of cells, we can run Dincta in an unsupervised way. It can infer the number of new cell types and annotate the cells into its correspond cell type, and do data integration keeping cells from different cell type separate while removing the batch effects to mix cells in the same cell type. Dincta is simple, accurate and efficient to integrate data, which keeps the cell type information preserved while removes the batch effects, and infers the known or novel cell types of cells.

## 1 Introduction

As the sequencing technology updates rapidly, we can sequence thousands of the cells in the gene expression levels in a short time. Now, there are huge mounts of single cell RNA-seq data generated and need to be analyzed. However, there are obstacles which hinder to analyze the scRNA-seq directly. The batch effects in the data which come from different samples, technologies or platforms hide the true biological interest variables. The researchers endeavor to solve this problem, and there emerges a bunch of tools for this task, such as MNN Correct (Lun et al., 2016), Seurat multiCCA (Butler et al., 2018), Scanorama (Hie et al., 2019), BBKNN (Polanski et al., 2019), Harmony (Korsunsky et al., 2018) and so on.

Harmony is one of the impressive tools to remove batch effects. It first projects the scRNA-seq raw counts data into a low dimension space (e.g. PCA). Secondly, it uses batch membership and cluster membership independent penalty to modify the soft *k*-means objective, which produces the soft cluster assignments. Third, it uses the the soft cluster assignments to do the linear correction of embedding, such that each cluster contains cells from different batch match to the batch frequency of the whole data. It iterates until the cluster assignments get stable.

There is a problem in Harmony. It uses a KL divergence loss to make that the batch membership is independent of the cluster membership. In the case that some cell type only occurs in part of samples (batches), this will case a problem. Since Harmony encourages that each cluster (subtype) consists of all batches. But in this case, the cluster should only contain cells in the batch which it contains the cell type. Harmony has a trend over correcting the batch effects.

If we know the cell type information of cells, this problem can be solved. We can constraint the penalty which only mixes cells in one cell type from different batches which contain the cell type, and do not mix cells from different cell types. That is the essential idea of Dincta. Here, we must to determine the cell type of cells. Note that in the k-means, each cluster is the surrogate of the cell types. As we reveal latter, the cell type assignment and cluster assignment can connect to each other via a probability relationship. The number of cell types can be determined by the rank of the cluster assignment matrix and we can construct the cell type assignment from the cluster assignment. Once getting the cell type information of the cells, we can move the low dimensional embedding of the cells such the embedding of cells belong to the same cell type will close to each other even they come from different batches, and the embedding of cells belong to different cell types separate a lot. As the embeddings of cells move toward the correct position, we will get more accurate cluster assignments and cell type assignments. And with the enhanced cluster assignments and cell type assignments, it will result in a more successful data integration. This build a positive enhance loop to make the Dincta converge to the final stable solution, i.e. correct cell type assignments and the data integration which mixes cells in the same cell type and keeps cells from different cell types separate.

## 2 Results

### 2.1 Dincta Mechanism

Dincta starts from the inputs of low dimensional embeddings (usually PCA embeddings) and the cell type information of the cells which have been annotated. It first run the constrained soft *k*-means with the the batch mixing penalty and cell type preserved penalty, to get the cluster assignment probability for each cluster. In the initial state, the the cluster may be separated by the batches. With the batch mixing penalty, it will adjust the cluster assignment probability for each cell such that the batch frequency of cells in this cluster approximates the batch frequency of cells in the cell type which the cluster belongs to. With the the cell type penalty, Dincta will adjust the cluster assignments to match the cell type assignments. Once we get the cluster assignment matrix, Dincta will infer the known or novel cell type of cells, which is coded in the cell type assignment matrix. After we get the batch mixing and cell type preserved adjusted cluster assignment matrix, Dincta use it to correct the embedding, such that the embedding of cells belong to the same cell type move to each other with a step, but the cells with different cell types keep separate. With the moved embeddings, it rerun the next time soft *k*-means clustering with the batch mixing and cell type preserved penalty. And do the novel cell type inferring if need, cell type assignment updating, embedding correcting, until finally the cluster assignments and cell type assignments get stable. After that, there is an option whether to refine the cell type assignment or not. If yes, Dincta will compute the fitness between the cluster assignments and cell type assignments for each cell with unknown cell type initially. If the cluster assignments and cell type assignments of a cell are not match to each other, then Dincta will mark this cell with unknown cell type; if they match to each other, Dincta will mark that cell with known cell type. After that, Dincta re-iterate the clustering, inferring and embedding correcting procedures described above until getting the stable and matchable cluster assignments and cell type assignments.

### 2.2 Performance Metric

We use three metrics to assess the results of Dincta. The first is the LISI (local inverse Simpson’s Index) metric (Korsunsky et al., 2018). It consists of cLISI (cell-type LISI) and iLISI (integration LISI). The cLISI is defined for each cell, it roughly measures the number of cell types in the local neighborhood of this cell. In the ideal case, we expect that cLISI approaches 1. The iLISI is also defined for each cell, it roughly measures the number of batches (samples, datasets, techonologies) in a neighborhood of this cell. If there are two batches, and each batch consists of the same cell types, we expect the the iLISI approaches 2.

The second metric is the match score (see in Methods **match score**) between the cluster assignments and the cell type assignments of cells. It was defined by the KL divergence of the cell type assignments and the cell type assignments which are transformed from the cluster assignments. The lower the value of match score, the fitness is better between the cluster assignments and cell type assignments.

The third metric is the inferring accuracy (see in Methods Algorithm 8). And there are two types of inferring accuracy. One type is that the cell type assignments come from the the cluster assignments directly (see Methods Algorithm 7) and comparing this cell type assignments with the golden cell type assignments of cells. The accuracy is determined by that the number of cells whose cell type were correctly inferred divided by the total number of cells (see Methods Algorithm 8). In the following tables, we denote this kind of inferring accuracy by the method name with suffix (with *R*). The another one is that cell type assignments come from the Dincta infer function (Algorithm 4) and comparing this cell type assignments with the golden cell type assignments of cells. In the ideal case, the inferring accuracy should approach 1.

In the all experiments below, we use the cell type annotated in Korsunsky et al. (2018) as golden labels of cells.

Besides the three metrics above, we also plot the density of the cLISI and iLISI to visually sense the behaviors of the algorithms. In order to visually check the results of cell type preserving and batch-diverse mixing, we plot the UMAP of the PCA embedding, Harmony embedding and Dincta embedding.

### 2.3 Quantifying Performance in Cell-Line Data

We first run Dincta on the cell-line data to check that it can do the correct jobs. The cell-line data consists of three samples. The first two samples: Jurkat and T293 consist of cells with the pure Jurkat cell type and pure T293 cell type, respectively. The third sample ‘half’ consists of cells with 50/50 mixing of two kinds of cell types: Jurkat and T293. And the cell type of cells can be labeled Jurkat or T293 unambiguously.

After pooling all the cells together, we performed a joint analysis. Before integration, cells group primarily by dataset (see Figure 1 (row at PCA), Table 1 (row at PCA) and Figure 2)

**Table 1:**
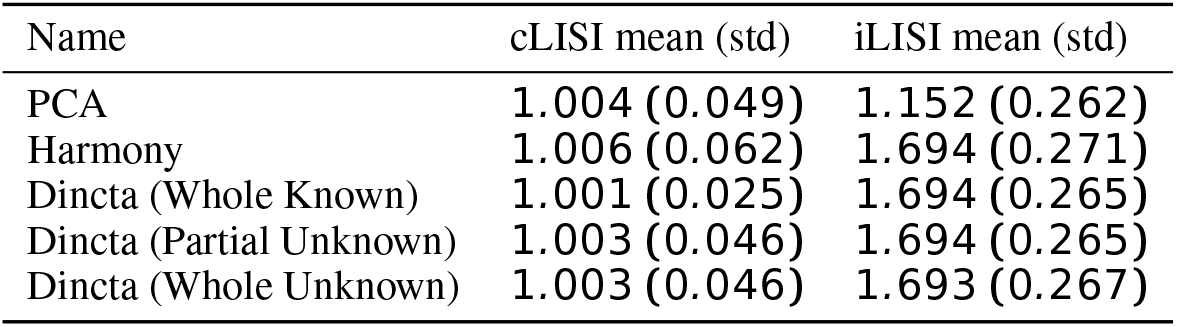
LISI Scores of Cell-Line Data

| Name | cLISI mean (std) | iLISI mean (std) |
| --- | --- | --- |
| PCA | 1.004 (0.049) | 1.152 (0.262) |
| Harmony | 1.006 (0.062) | 1.694 (0.271) |
| Dincta (Whole Known) | 1.001 (0.025) | 1.694 (0.265) |
| Dincta (Partial Unknown) | 1.003 (0.046) | 1.694 (0.265) |
| Dincta (Whole Unknown) | 1.003 (0.046) | 1.693 (0.267) |

**Figure 1:**
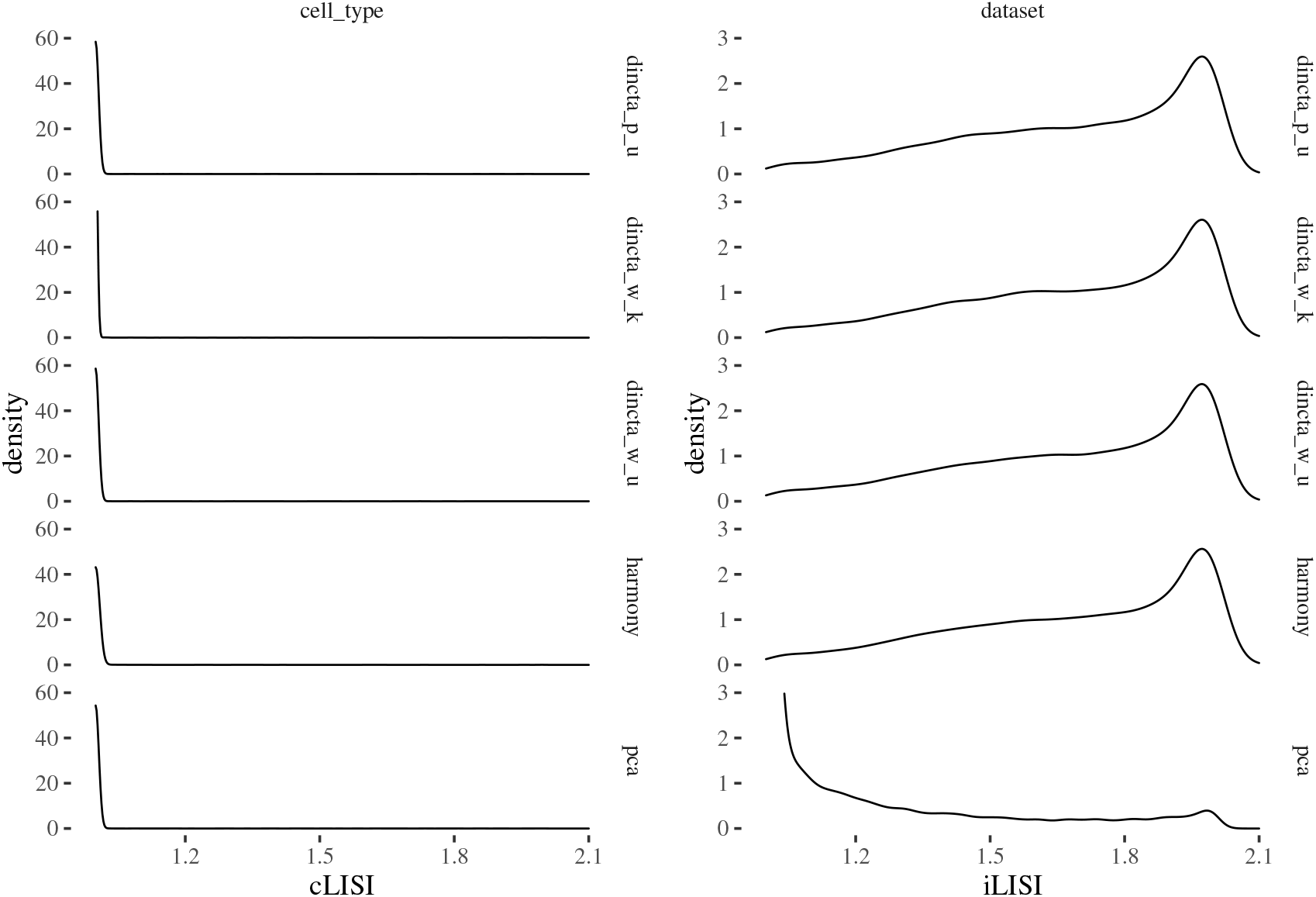
LISI density of cell-line data

**Figure 2:**
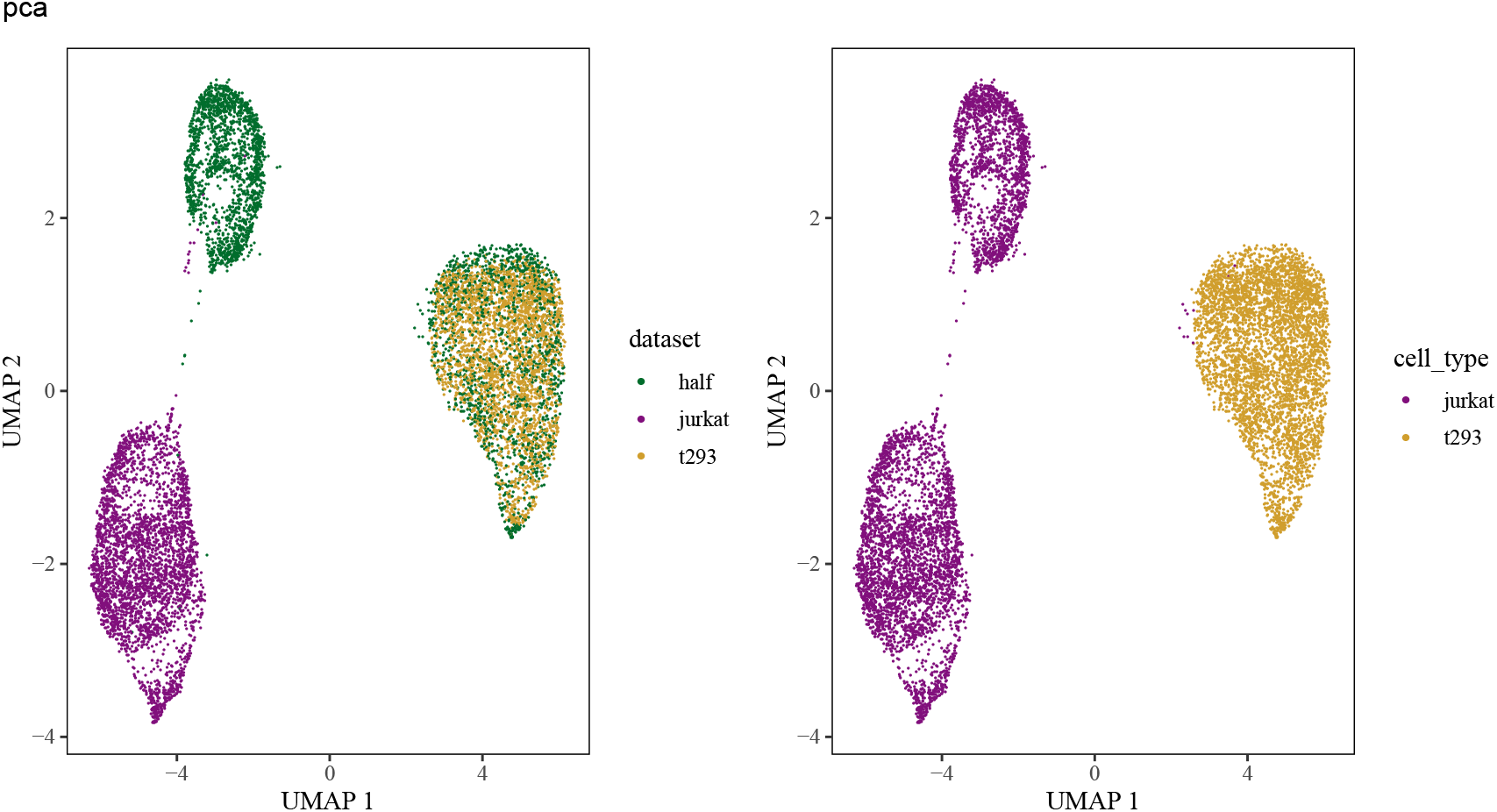
PCA results of cell-line data.

We perform three experiments on the cell-line data. In the first experiment, we use the whole cell annotation of cells annotated in the Korsunsky et al. (2018). We feed these cell type information and the PCA embedding to the Dincta. After Dincta, we compare the integration results coming from the Harmony (Figure 3) with Dincta (Figure 4. The UMAP figures of Dincta and Harmony show that both algorithms get the similar results. Once zooming in the cell type UMAP figure of Harmony, we can find several misclassified cells in the center of cells with Jurkat cell type. Dincta also misclassified few cells but the number of misclassified cells is less than Harmony’s. The inferring accuracy table (Table 3 (rows at Dincta (Whole Known with *R*) and Harmony (with *R*)) also verifies this observation. We see that Dincta perform better than Harmony in the match score as show in table 2 (rows at Harmony and Dincta (Whole Known)). From the LISI results (Table 1 (rows at Dincta (Whole Known with *R*) and Harmony) and Figure 1 (row as dincta_w_k)), Dincta has lower mean value of cLISI which show that it preserves the cell type separate better than Harmony while keeps the similar iLISI values. This indicates that the Harmony over corrects the batch effects a little and mixes a few cells with different cell types.

**Table 2:** Match Score between Cell Type Assignment and Cluster Assignment of Cell-Line Data

| Name | Match Score |
| --- | --- |
| Harmony | 131.8 |
| Dincta (Whole Known) | 35.5 |
| Dincta (Partial Unknown) | 30.6 |
| Dincta (Whole Unknown) | 31.0 |

**Table 3:** Inferring Accuracy of Cell-Line Data

| Name | accuracy | number of infer correct cells | total number of cells |
| --- | --- | --- | --- |
| Harmony (with $R$) | 0.9942 | 9423 | 9478 |
| Dincta (Whole Known with $R$) | 0.9994 | 9472 | 9478 |
| Dincta (Partial Unknown ) | 0.9979 | 9458 | 9478 |
| Dincta (Partial Unknown with $R$) | 0.9979 | 9458 | 9478 |
| Dincta (Whole Unknown) | 0.9979 | 9458 | 9478 |
| Dincta (Whole Unknown with $R$) | 0.9978 | 9457 | 9478 |

**Figure 3:**
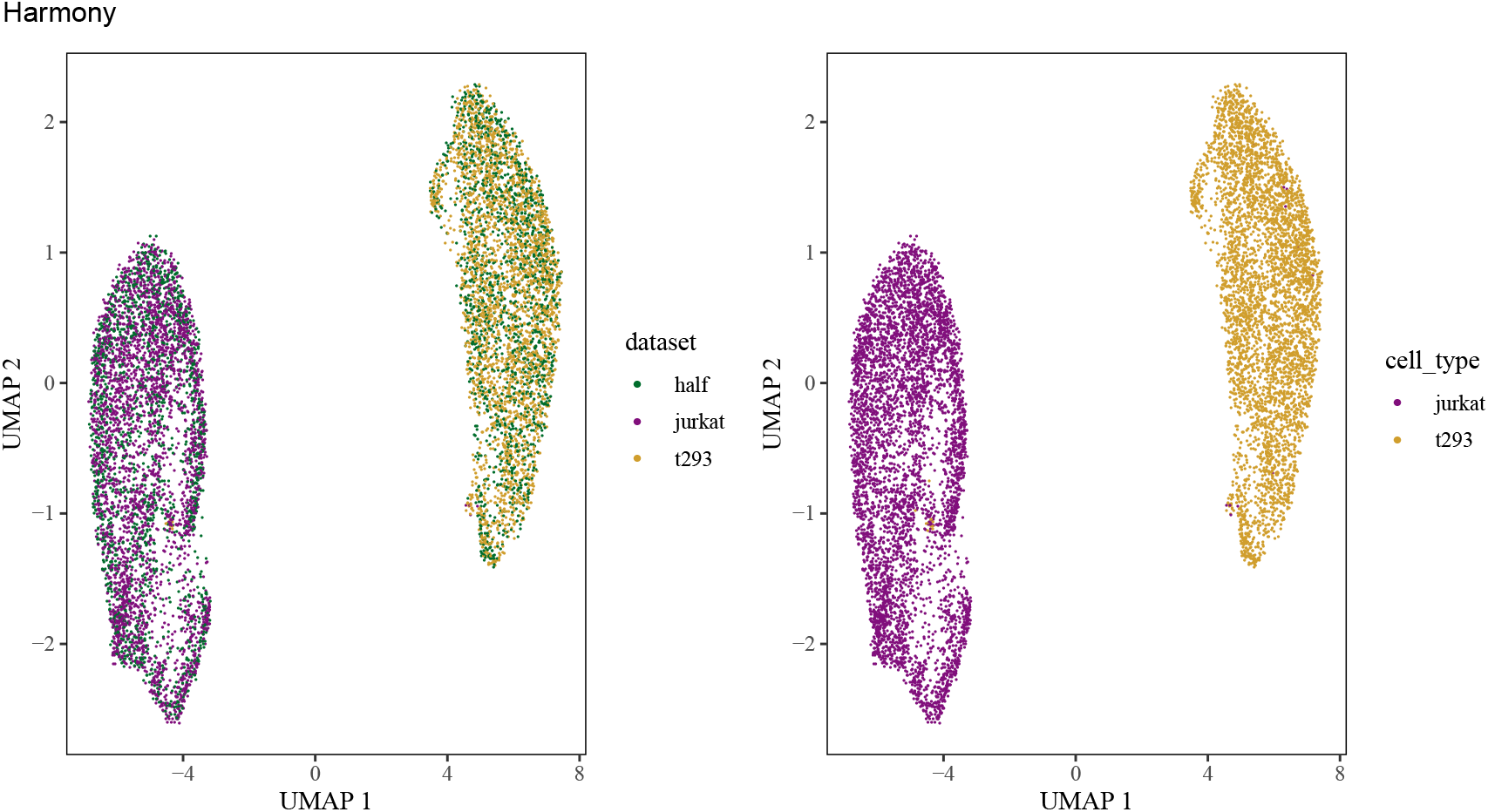
Harmony results of cell-line data.

**Figure 4:**
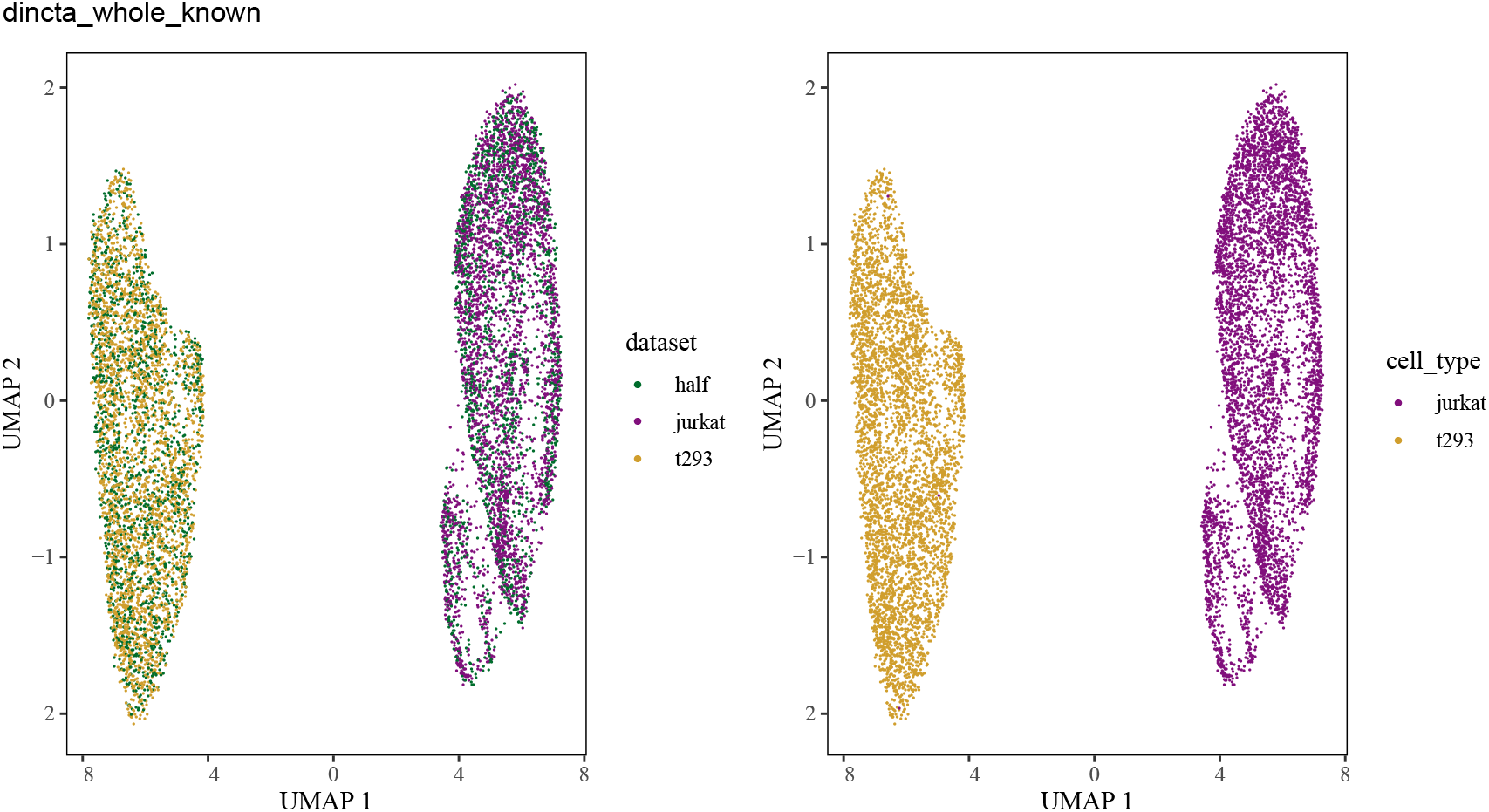
Dincta (with whole cell type information as input) results of cell-line data.

In the second experiment, we mask the cell type information of cells in half and t293 dataset to the “unknown” cell type, only provide the cell type information of cells in jurkat dataset. We feed the partial cell type information (fake_cell_type) and the PCA embedding to the Dincta. After Dincta, we plot the UMAP of Dincta embedding as showed in Figure 5). We see that Dincta performs similarly as we feed all the cell type information. Dincta infers the the masked “T293” cell type as the “new_cell_type_1”. And it assigns the cell type of cells with the masked “unknown” cell type initially almost correctly, which can be checked from the Table 3 (rows at Dincta (Partial Unknown) and Dincta (Partial Unknown with *R*)). From Table 1 (row at Dincta (Partial Unknown)) and Figure 1) (row at Dincta_p_u), we find Dincta also performs a litter better than results of the Harmony. This experiment reveals a road to do the cell type annotation in practice. First, we run a cluster algorithm (e.g. DinctaClassification) on one sample, and use the marker genes to determine the cell type of cells. Then when a new sample comes, we can run the Dincta to infer the known or novel cell type of cells in the new sample.

**Figure 5:**
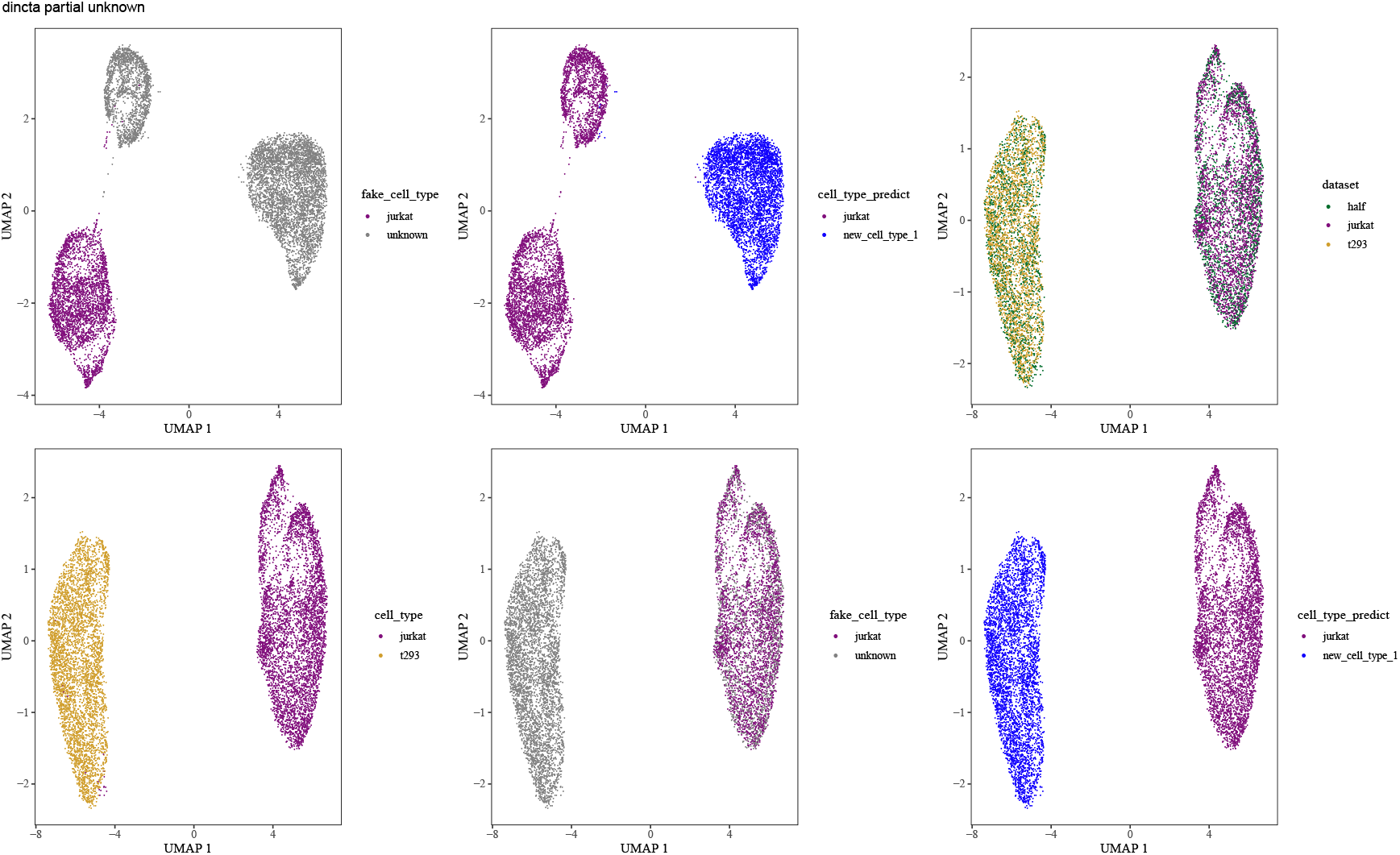
Dincta (with partial cell type information) results of cell-line data. The top left and top middle figures are the UMAP of the PCA embedding, colored by the partial cell types (fake_cell_type) of cells which is cell type information input to Dincta and colored by the inferred cell type information (cell_type_predict), respectively. The rest four figures are the UMAP of the Dincta embedding colored by dataset, cell_type, fake_cell_type and cell_type_predict, respectively.

In the third experiment, we mask the cell type of all cells to the “unknown” cell type. After Dincta, it infers that there are 2 cell types in the data. From the figure 6, we can easily find that “new_cell_type_1” is the “T293” cell type, and “new_cell_type_2” is the “Jurkat” cell type. Dincta achieves almost the same cell type assignments as the cell type of cells which were annotated separately in each dataset by the marker genes. Comparing the inferring accuracy, we see that Dincta achieves very high Inferring Accuracy with only a little performance drop (Table 3(rows at Dincta (Whole Unknown) and Dincta (Whole Unknown with *R*))). From Table 2 (row at Dincta (Whole Unnown)), we see that the cluster assignments and cell type assignments also fit. This experiment shows that Dincta is robust even under all the cell type information is masked out.

**Figure 6:**
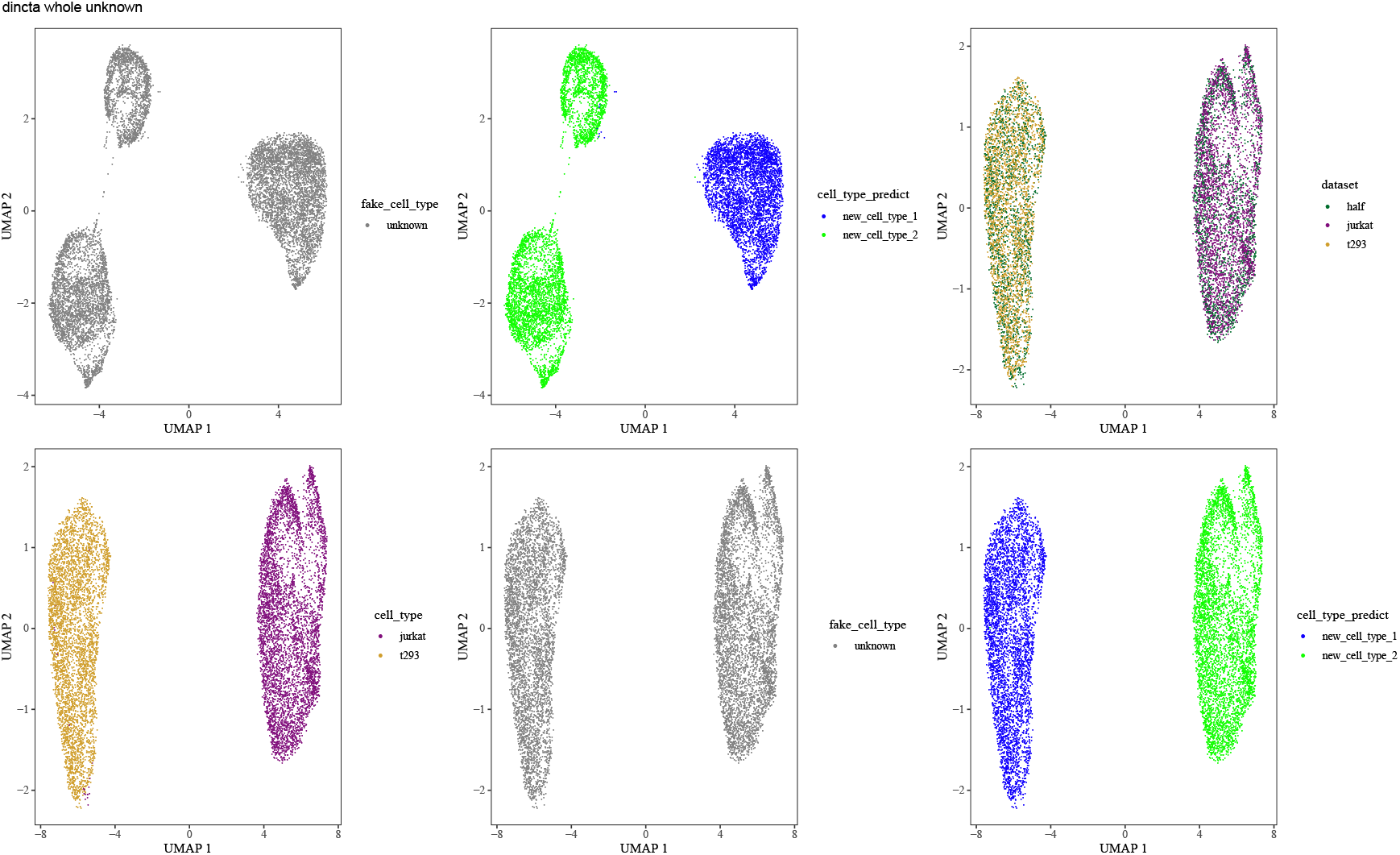
Dincta (with no cell type information) results of cell-line data. The top left and top middle figures are the UMAP of the PCA embedding, colored by the no cell type information (unknown) (fake_cell_type) which is cell type information input to the Dincta and colored by the inferred cell type information (cell_type_predict), respectively. The rest four figures is the UMAP of the Dincta embedding colored by dataset, cell_type, fake_cell_type and cell_type_predict, respectively.

### 2.4 Quantifying Performance in PBMC data

To assess how Dincta might perform under more challenging scenarios, we gathered three dataset of human PBMCs, each assayed with the Chromium 10X platform but prepared with different protocols: 3′ and *v1* (3pV1), 3′ end *v2* (3pV2) and 5′ end (5p) chemistries. To make the task harder, we remove the cells with “mono” cell type in protocol 5p, remove the cells with “tcells” cell type in protocol 3pV1 and remove the cells with “bcells” cell type in protocol 3pV2. The composition of cell types for the three datasets tabulates in Table 4

**Table 4:**
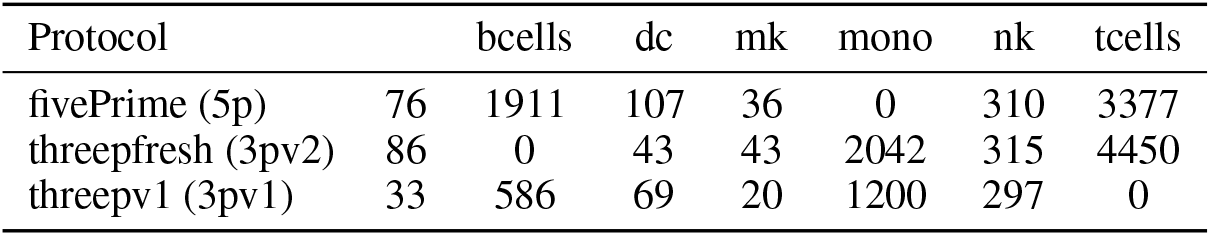
Cell Type Composition of PBMCs

After pooling all the cells together, we performed a joint analysis. Before integration, cells group primarily by dataset (see Figure 7 (row at PCA), Table 5 (row at PCA) and Figure 8).

**Table 5:** LISI Scores of PBMC data

| Name | cLISI mean (std) | iLISI mean (std) |
| --- | --- | --- |
| PCA | 1.017 (0.112) | 1.015 (0.110) |
| Harmony | 1.042 (0.161) | 1.682 (0.345) |
| Dincta (Whole Known) | 1.014 (0.103) | 1.640 (0.389) |
| Dincta (Partial Unknown) | 1.017 (0.107) | 1.662 (0.378) |
| Dincta (Whole Unknown) | 1.034 (0.141) | 1.646 (0.355) |

**Figure 7:**
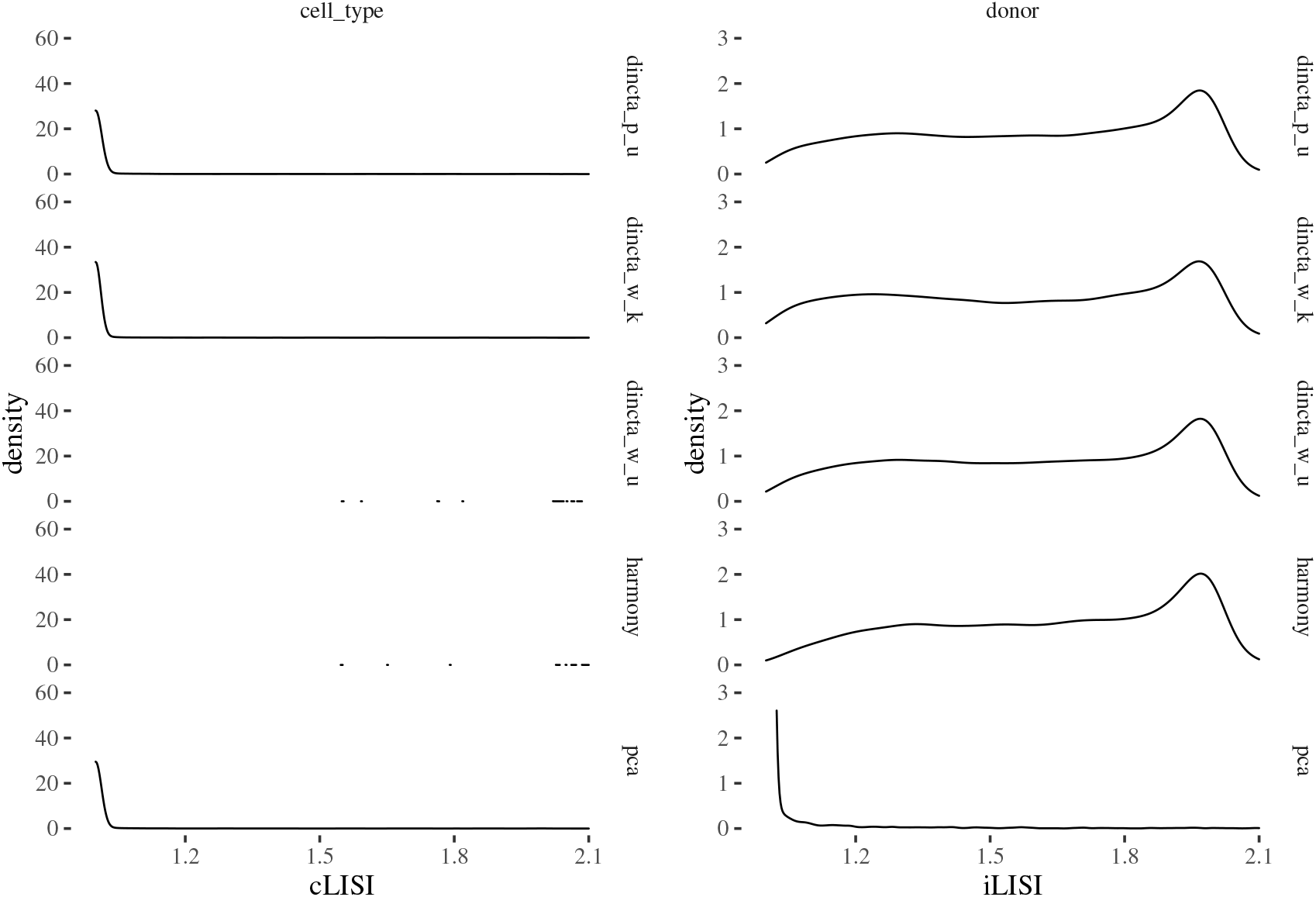
LISI of PBMC data.

**Figure 8:**
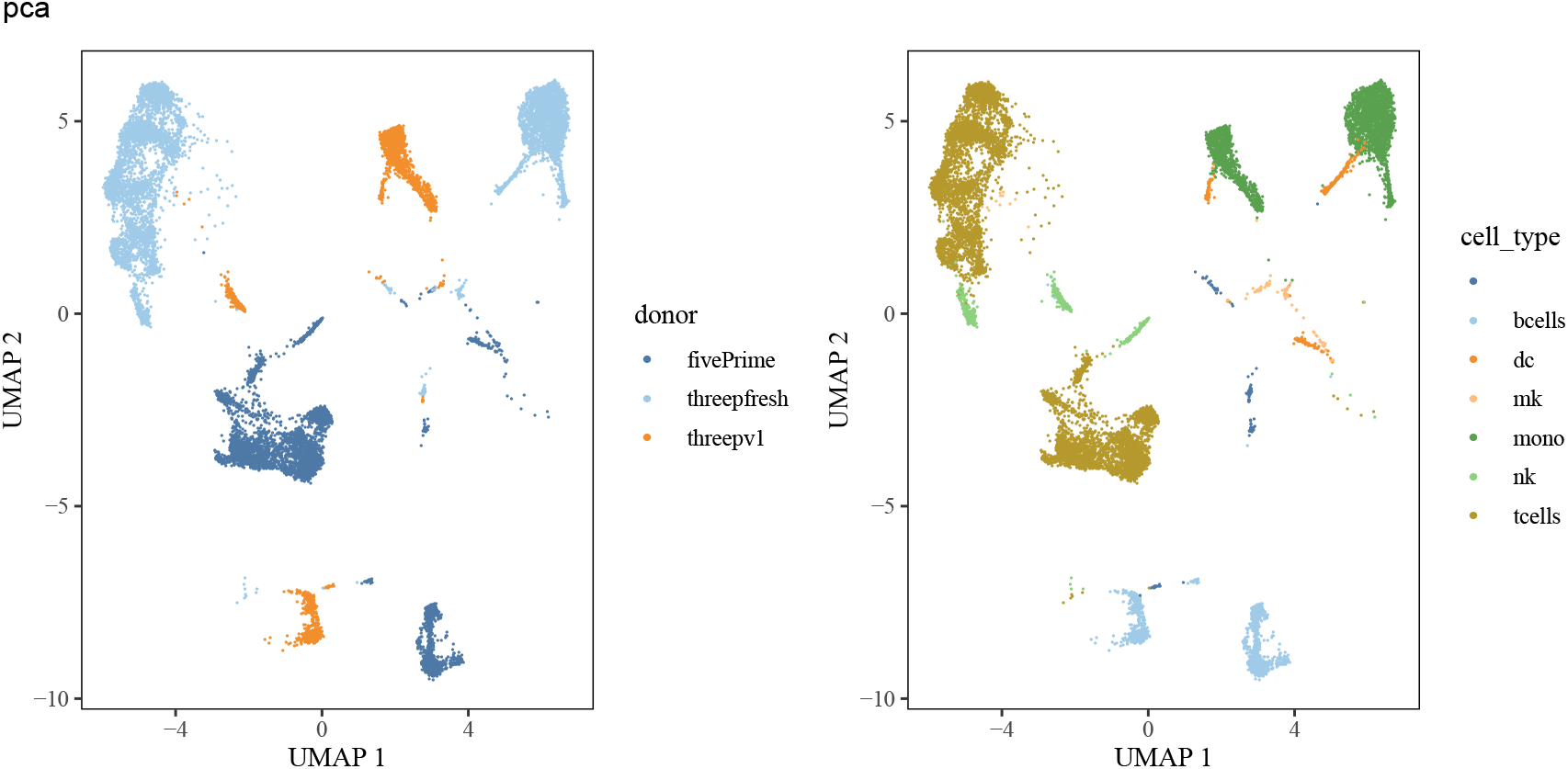
PCA results of PBMC data.

We perform three experiments on the PBMCs data. In the first experiment, we use the whole cell annotation of cells annotated in the Korsunsky et al. (2018). We feed these cell type information and the PCA embedding to the Dincta. After Dincta, we compare the integration results coming from the Harmony (Figure 9) with Dincta (Figure 10). Once zooming in Figure 9, we find that in the cluster of “bcells”, there are some cells from protocol 3pv2. This should not happen, since the protocol 3pv2 don’t own cells with “bcells” cell type. The similar phenomena occur in the clusters of cell type “mono” and “tcells”. It means that Harmony over corrects the batch effects and mixes a small part of cells with different cell types. We zoom in Figure 10, Dincta does a better job. In the cluster of “bcells”, it consists of cells from protocols 5p and 3pv1 majorly, and only few cells comes from protocol 3pv2. The similar phenomena also occur in the cluster of cells with “mono” and “tcells” cell type. This shows that Dincta mixes the cells with the same cell type from different batches (protocols), while keeps the cells with different cell types separate. This can be also verified from Table 5(rows at Harmony and Dincta (Whole Known)), Figure 7(rows at dincta_w_k and harmony)), Table 2 (rows at Harmony and Dincta (Whole Known)) and Table 7 (rows at Dincta (Whole Known with *R*) and Harmony (with *R*)).

**Figure 9:**
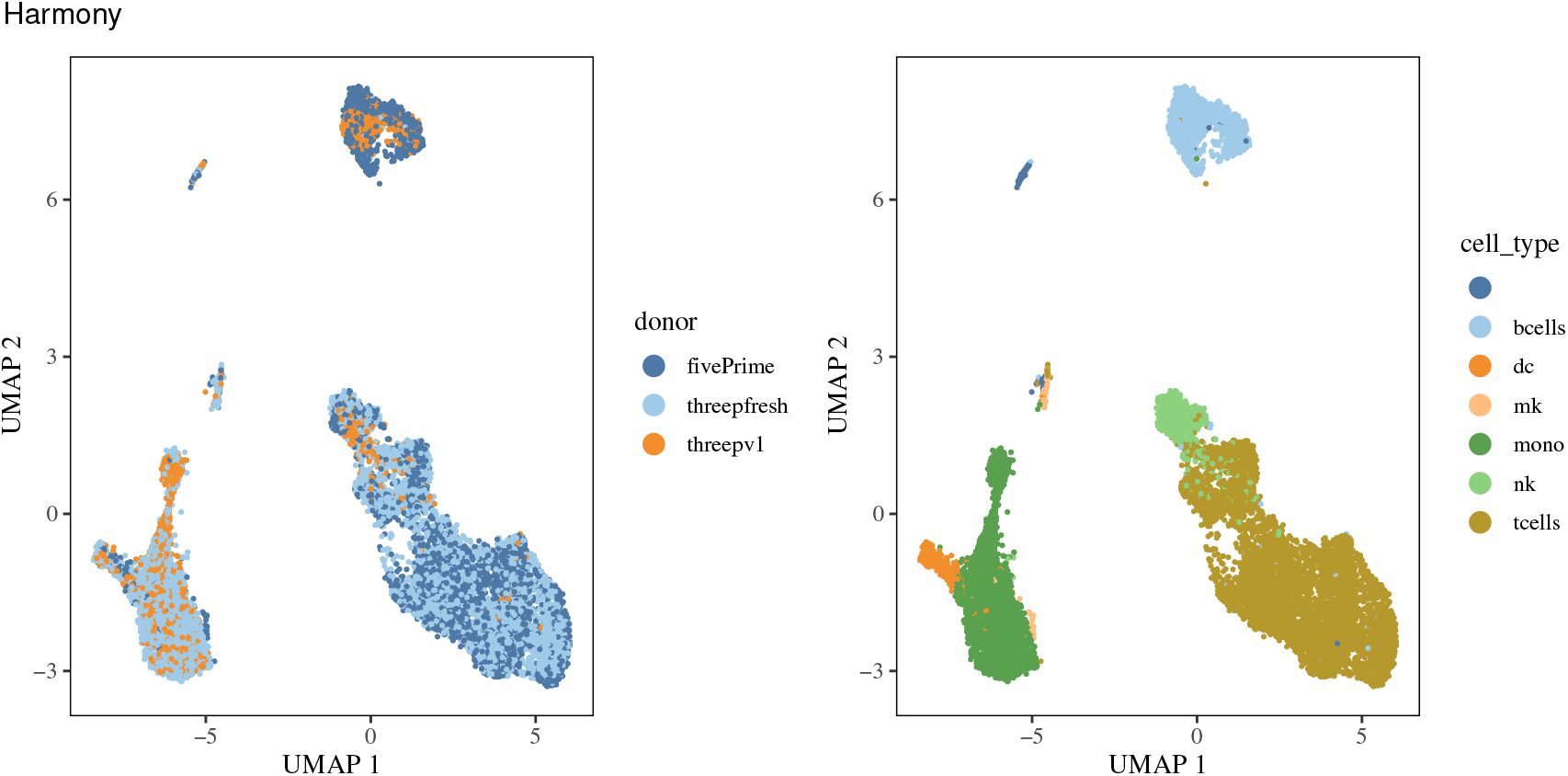
Harmony results of PBMC data.

**Figure 10:**
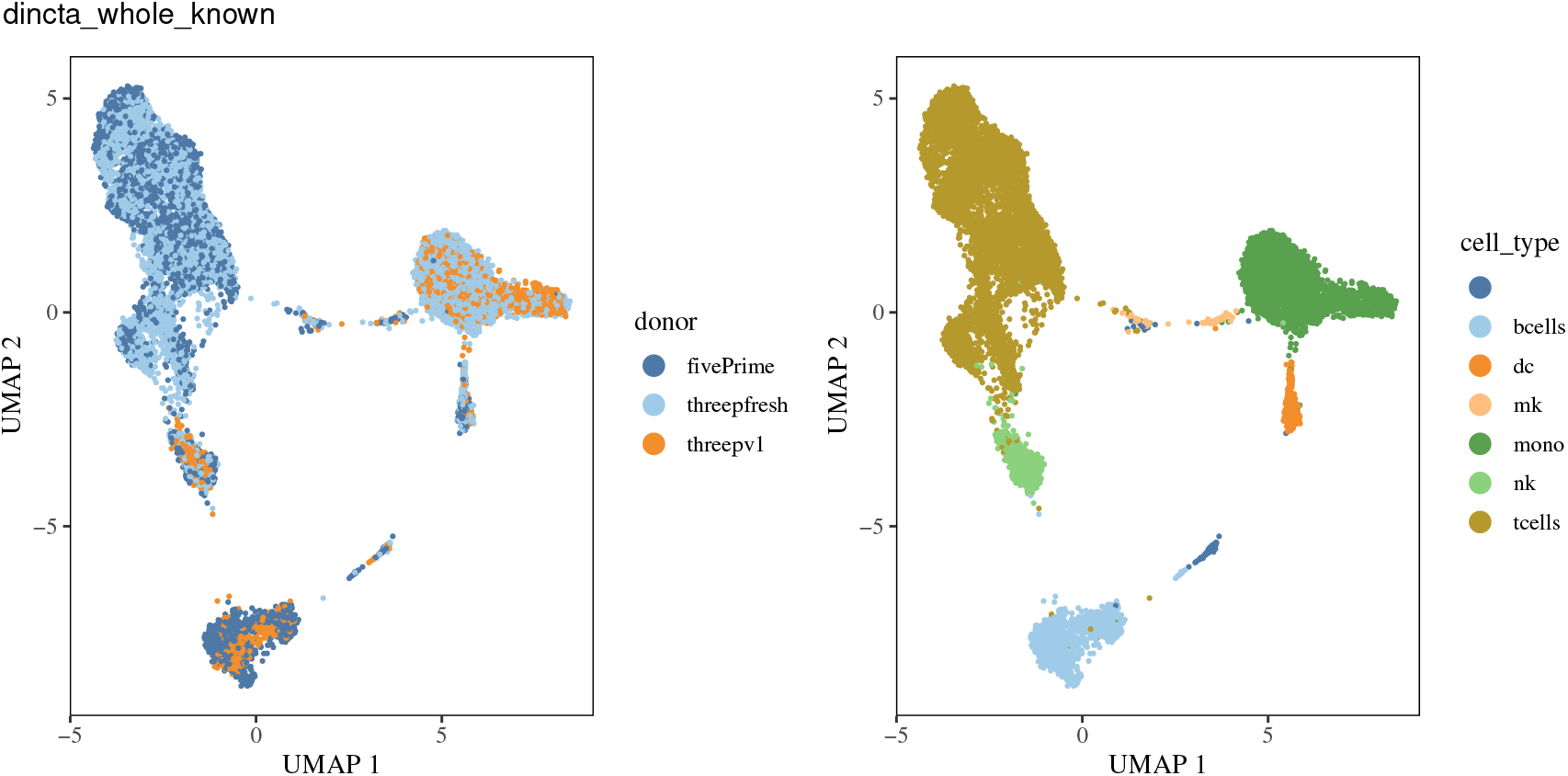
Dincta (with whole cell type information as input) results of PBMC data.

In the second experiment, we mask the the cell type information of cells measured with **5’** end (5p) chemistries protocol to the “unknown” cell type. And we also mask the cell type “mono” for all cells as “unknown” cell type. We feed the partial cell type information (fake_cell_type) and the PCA embedding to the Dincta. From the Figure 11), we see that Dincta performs similarly as we feed all the cell type information. Dincta infers the the masked “mono” cell type as the two subtypes “new_cell_type_1” and “new_cell_type_2”. And it assigns the cell type of cells with the masked “unknown” cell type initially with high accuracy, which can be checked from the Table 7 (rows at Dincta (Partial Unknown) and Dincta (Partial Unknown with *R*)). From Table 5 (row at Dincta (Partial Unknown)) and Figure 7) (row at Dincta_p_u), we find Dincta also performs better than Harmony.

**Figure 11:**
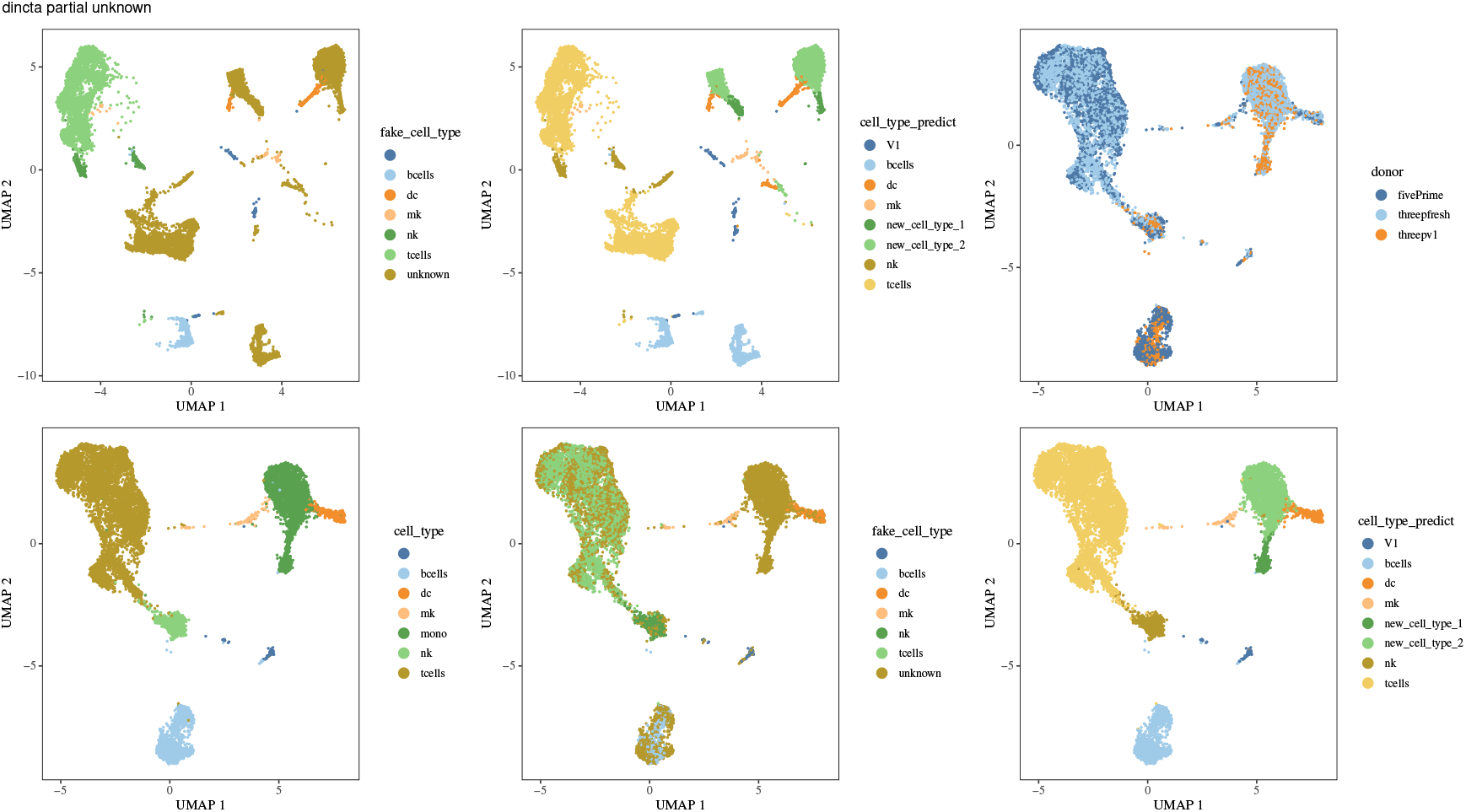
Dincta (with partial cell type information) results of PBMC data. The top left and top middle figures are the UMAP of the PCA embedding, colored by the partial cell types (fake_cell_type) of cells which is cell type information input to Dincta and colored by the inferred cell type information (cell_type_predict), respectively. The rest four figures are the UMAP of the Dincta embedding colored by donor, cell_type, fake_cell_type and cell_type_predict, respectively.

In the third experiment, we mask the cell type of all cells to the “unknown” cell type. After Dincta, it infers that there are 9 cell types in the data. As we see in the Figure 12, Dincta achieves similar cell type assignment results as the cell type of cells which were annotated separately in each dataset by the marker genes. There is some mismatch between the golden labels and the predicted cell types. Dincta annotates the the cells with “dc” cell type and part of cells with “mono” cell type to the single “new_cell_type_9”. And part of cells with “mono” cell type was annotated to the “new_cell_type_3”. Also the cells with the “tcells” cell type are annotated with subtypes “new_cell_type_4”, “new_cell_type_5” and “new_cell_type_7”. So there still has the improve space of Dincta. Although it has these mistakes, Dincta almost keeps the cells with different inferred new cell types separate. From the cluster of cells with “bcells” cell type, there was almost no cells from the protocol 3pv2, which is better than the results of Harmony. The similar phenomena occur in the clusters of cells with “tcells” and “mono” cell type. From Table 7(rows at Dincta (Whole Unknown) and Dincta (Whole Unknown with *R*)), Table 6 (row at Dincta (Whole Unnown)), Dincta achieves a little better results than Harmony’s.

**Table 6:** Match Score between Cell Type Assignment and Cluster Assignment of PBMCs

| Name | Match Score |
| --- | --- |
| Harmony | 2188.6 |
| Dincta (Whole Known) | 84.4 |
| Dincta (Partial Unknown) | 650.3 |
| Dincta (Whole Unknown) | 1889.5 |

**Table 7:** Inferring Accuracy of PBMCs

| Name | accuracy | number of infer correct cells | total number of cells |
| --- | --- | --- | --- |
| Harmony (with $R$) | 0.8964 | 12962 | 14460 |
| Dincta (Whole Known with $R$) | 0.9982 | 14434 | 14460 |
| Dincta (Partial Unknown ) | 0.9539 | 13794 | 14460 |
| Dincta (Partial Unknown with $R$) | 0.9049 | 13086 | 14460 |
| Dincta (Whole Unknown) | 0.9128 | 13200 | 14460 |
| Dincta (Whole Unknown with $R$) | 0.9019 | 13042 | 14460 |

**Figure 12:**
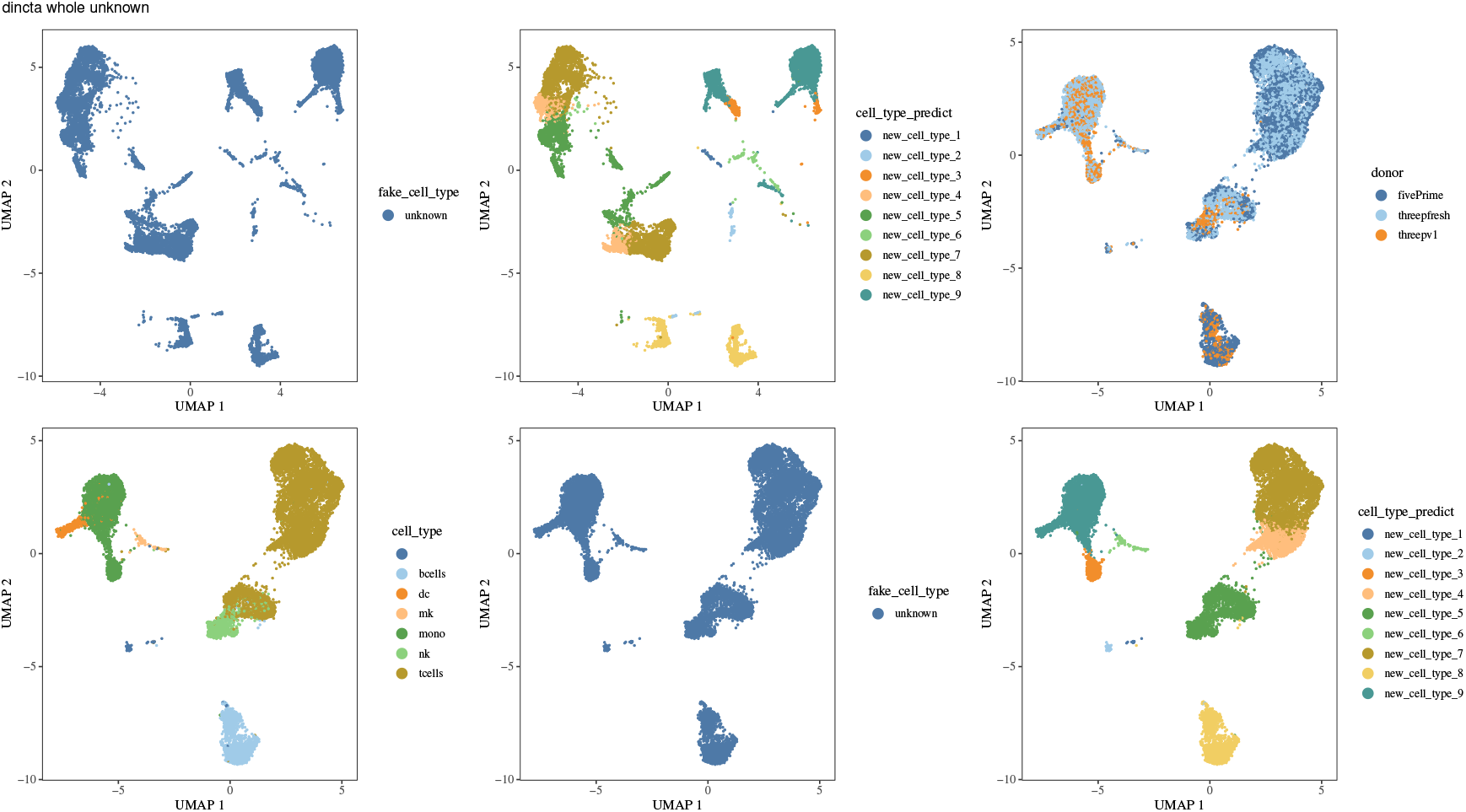
Dincta (with no cell type information) results of PBMC data. The top left and top middle figures are the UMAP of the PCA embedding, colored by the no cell type information (unknown) (fake_cell_type) which is cell type information input to the Dincta and colored by the inferred cell type information (cell_type_predict), respectively. The rest four figures is the UMAP of the Dincta embedding colored by donor, cell_type, fake_cell_type and cell_type_predict, respectively.

### 2.5 Quantifying Performance in Pancreas Islet Data

To assess Dincta on a more complex experimental design, we gathered human pancreatic islet cells from three independent studies as described in Korsunsky et al. (2018), each generated with a different technological platform. The batch effects result from both different technologies and the **36** donors used in these studies. A successful integration of these studies must be performed simultaneously over the variations from the different technologies and donors.

After pooling all the cells together, we performed a joint analysis. Before integration, cells group primarily by dataset (see Figure 13 (row at PCA), Table 8 (row at PCA) and Figure 14).

**Table 8:** LISI Scores of Pancreas Data

| Name | cLISI mean (std) | iLISI mean (std) (dataset) | iLISI mean (std) (donor) |
| --- | --- | --- | --- |
| PCA | 1.061 (0.213) | 1.028 (0.147) | 1.904 (1.216) |
| Harmony | 1.067 (0.233) | 1.832 (0.701) | 4.298 (2.082) |
| Dincta (Whole Known) | 1.059 (0.184) | 1.781 (0.676) | 4.158 (2.043) |
| Dincta (Partial Unknown) | 1.057 (0.174) | 1.794 (0.682) | 4.203 (2.068) |
| Dincta (Whole Unknown) | 1.068 (0.227) | 1.807 (0.692) | 4.218 (2.051) |

**Figure 13:**
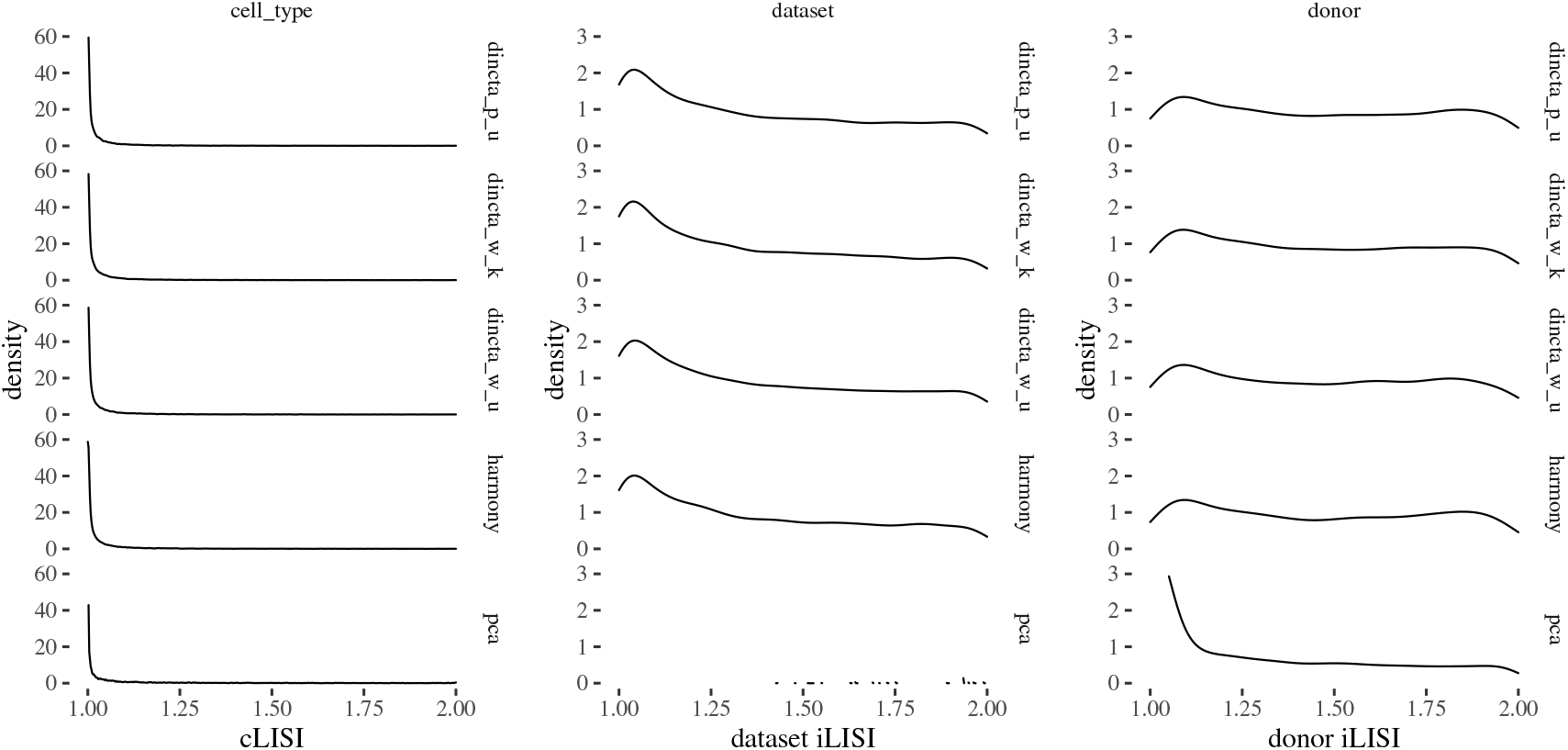
LISI of pancreas data

**Figure 14:**
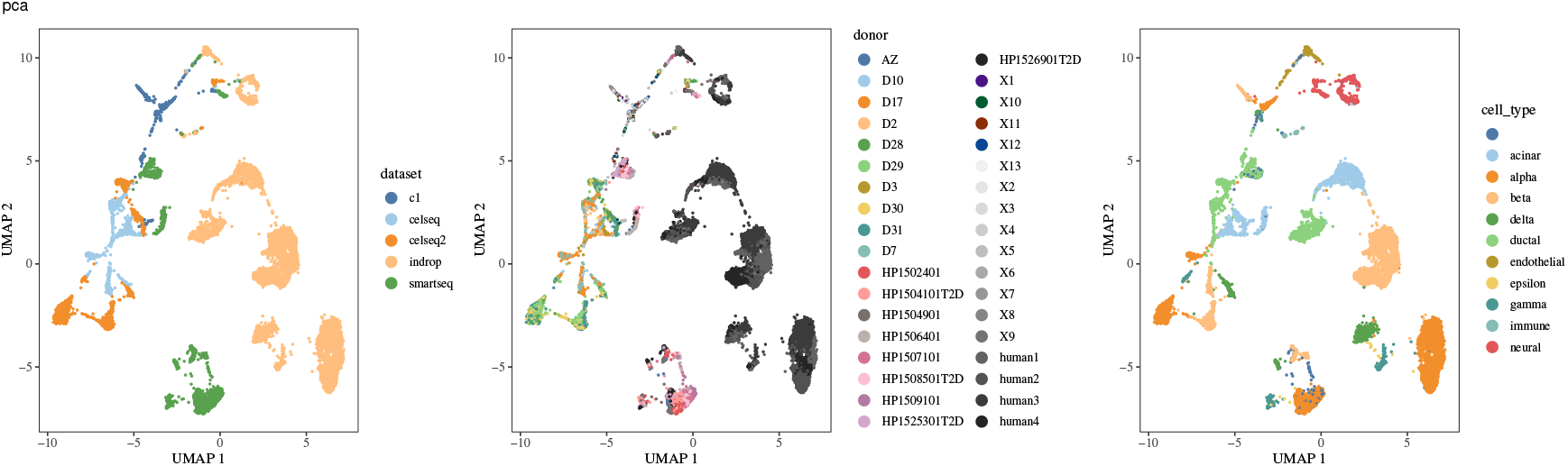
PCA results of pancreas data.

We perform three experiments on the pancreas islet data.

In the first experiment, we use the whole cell type information of cells. We feed the whole cell type information and the PCA embedding to the Dincta. After Dincta, we compare the integration results coming from the Harmony (Figure 15) with Dincta (Figure 16). The cluster of cells with “neural” cell type consists of cells with almost the pure “neural” cell type in the UMAP of Dincta’s embedding, while the cluster of cells with “neural” cell type contains several cells with “endothelial” cell type in the UMAP of Harmony’s embedding. The similar phenomena also occur in the clusters of cells with other cell types. This shows that Dincta mixes the cells with the same cell type from different batches, while tries to keep the cells with different cell types separate. This can be also verified from Table 8(rows at Harmony and Dincta (Whole Known)), Figure 13(rows at dincta_w_k and harmony)), Table 9 (rows at Harmony and Dincta (Whole Known)) and Table 10 (rows at Dincta (Whole Known with *R*) and Harmony (with *R*)). In this experiment, Dincta does a better job than Harmony.

**Table 9:**
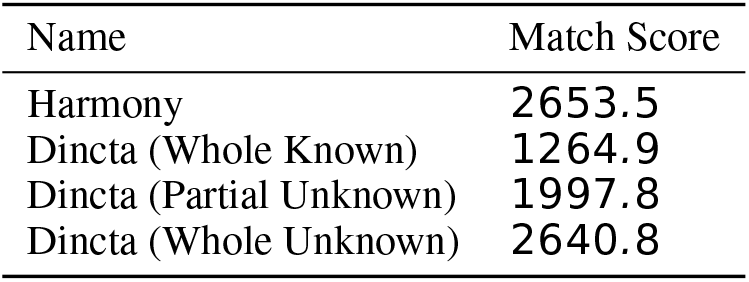
Match Score between Cell Type Assignment and Cluster Assignment of Pancreas Data

**Table 10:** Inferring Accuracy of Pancreas Data

| Name | accuracy | number of infer correct cells | total number of cells |
| --- | --- | --- | --- |
| Harmony (with $R$) | 0.9616 | 14180 | 14746 |
| Dincta (Whole Known with $R$) | 0.9775 | 14415 | 14746 |
| Dincta (Partial Unknown ) | 0.9710 | 14319 | 14746 |
| Dincta (Partial Unknown with $R$) | 0.9697 | 14300 | 14746 |
| Dincta (Whole Unknown) | 0.9622 | 14189 | 14746 |
| Dincta (Whole Unknown with $R$) | 0.9622 | 14189 | 14746 |

**Figure 15:**
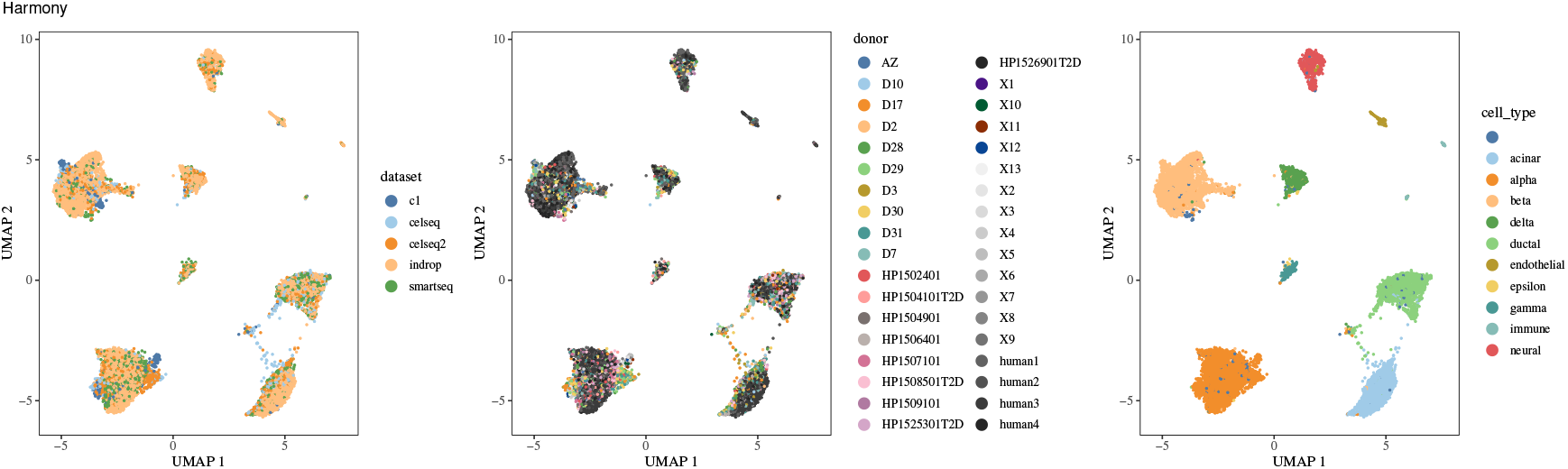
Harmony results of pancreas data.

**Figure 16:**
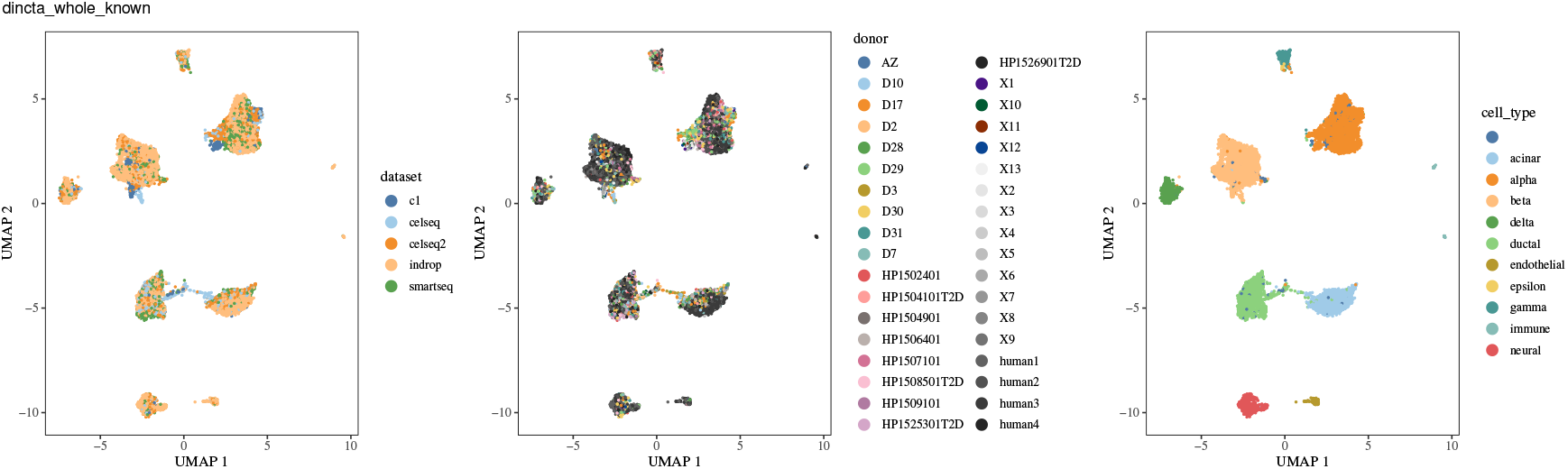
Dincta (with whole cell type information as input) results of pancreas data.

In the second experiment, we mask the the cell type information of cells measured with celseq2 to the “unknown” cell type. And we also mask the cell type “alpha” for all cells as “unknown” cell type. After Dincta, we plot the UMAP of Dincta embedding as showed in Figure 17). We see that Dincta performs similarly as we feed all the cell type information. Dincta infers the the masked “alpha” cell type as “new_cell_type_1”. And it assigns the cell type of cells with the masked “unknown” cell type initially with high accuracy, which can be checked from the Table 10 (rows at Dincta (Partial Unknown) and Dincta (Partial Unknown with *R*)). From Table 8 (row at Dincta (Partial Unknown)) and Figure 13) (row at Dincta_p_u), we find Dincta also performs a little better than results of the Harmony.

**Figure 17:**
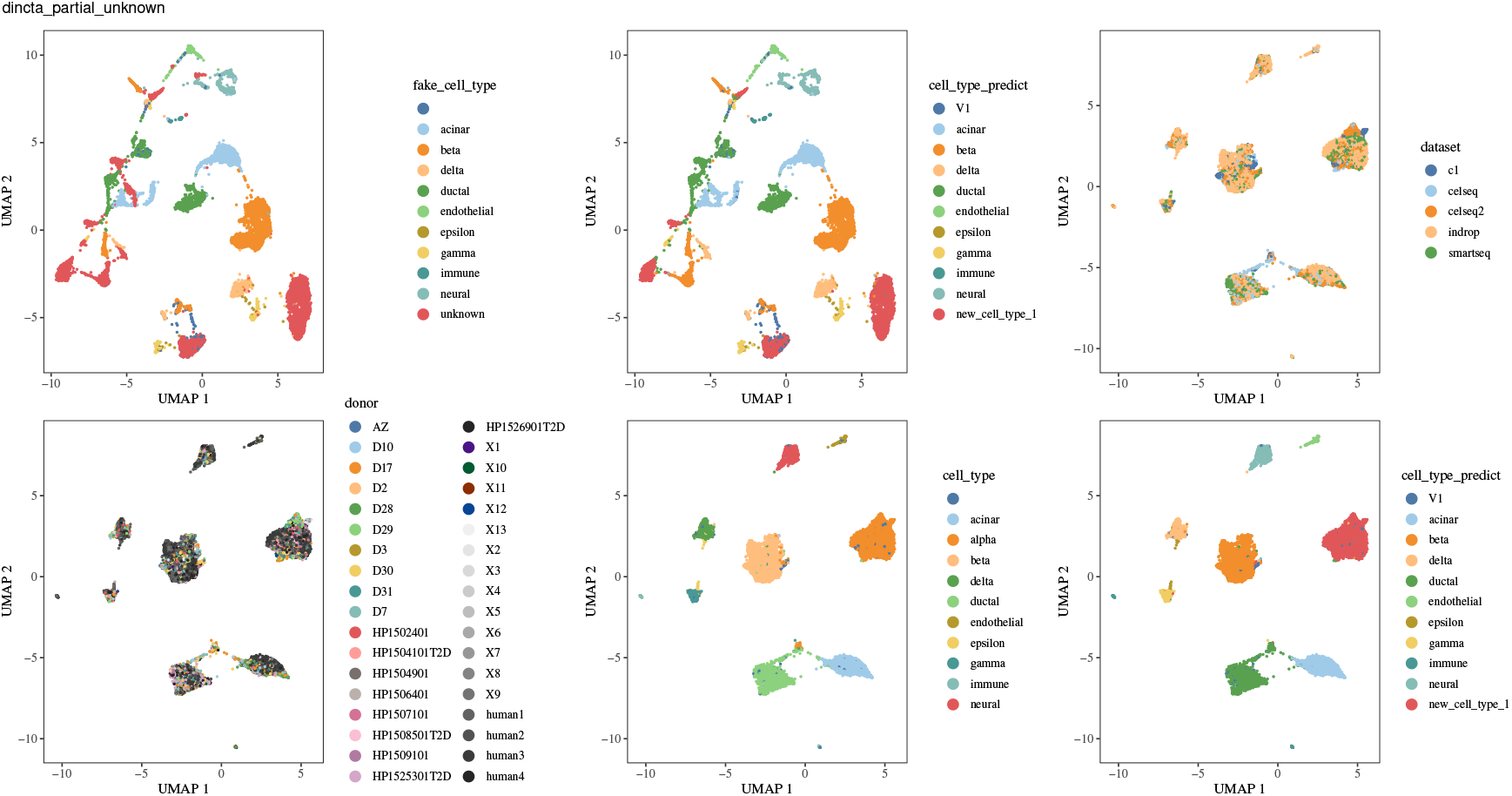
Dincta (with partial cell type information) results of pancreas data. The top left and top middle figures are the UMAP of the PCA embedding, colored by the partial cell types (fake_cell_type) of cells which is cell type information input to Dincta and colored by the inferred cell type information (cell_type_predict), respectively. The rest four figures are the UMAP of the Dincta embedding colored by dataset, cell_type, fake_cell_type and cell_type_predict, respectively.

In the third experiment, to make the task more harder, we mask the cell type of all cells to the “unknown” cell type. After Dincta, it infers that there are 10 cell types in the data. As we see in the Figure 18, Dincta achieves similar cell type assignment results as the cell type of cells which were annotated separately in each dataset by the marker genes. Dincta almost keeps the cells with different inferred new cell types separate. From Table 9 (row at Dincta (Whole Unnown)), we see that the cluster assignments and cell type assignments also fit to each other better than Harmony’s results. From Table 10 (rows at Dincta (Whole Unknown) and Dincta (Whole Unknown with *R*)), Dincta achieves a little better results than Harmony’s. This experiment shows that Dincta is robust even under all the cell type information is masked out.

**Figure 18:**
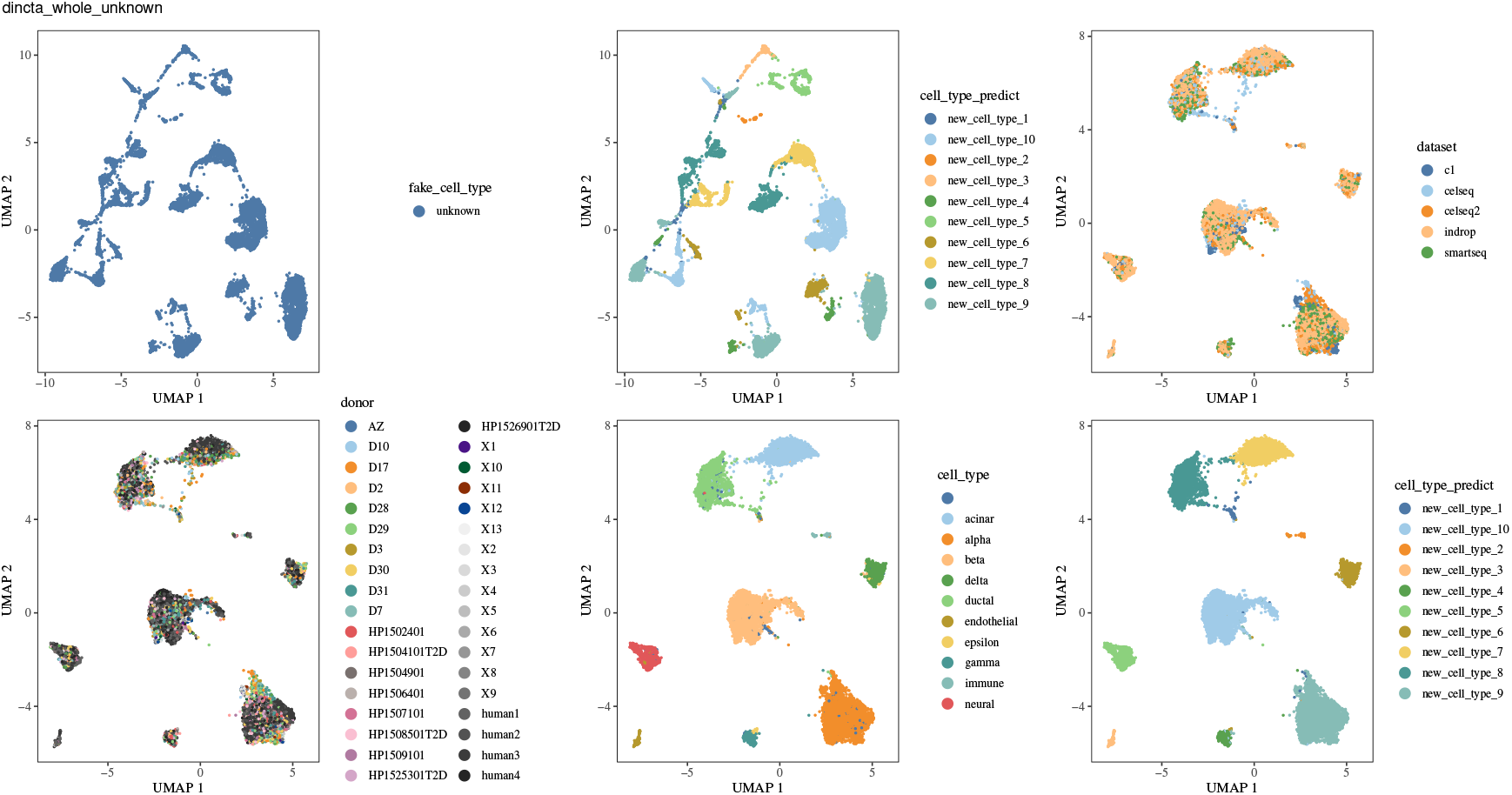
Dincta (with no cell type information) results of pancreas data. The top left and top middle figures are the UMAP of the PCA embedding, colored by the no cell type information (unknown) (fake_cell_type) which is cell type information input to the Dincta and colored by the inferred cell type information (cell_type_predict), respectively. The rest four figures is the UMAP of the Dincta embedding colored by dataset, cell_type, fake_cell_type and cell_type_predict, respectively.

## 3 Discussion

Dincta builds a unify framework to do the data integration and cell type annotation simultaneously. We can input all or partial or none of the cell type information of cells which are annotated ahead of running Dincta. Dincta can use these cell type information to infer known or novel cell type of cells, cluster cells in the same cell type together while keeping cells from different cell types separate.

When there is only one batch, this problem reduce to the cell type inference and assignment problem with no need to do data integration. We implemented this simple case in the function “DinctaClassification” of Dincta Package.

Currently, Dincta is based on the frequency clustering problem essentially. In the future, it is worth to generalize the algorithm to case which use the count values of mRNA.

## 4 Methods

We now begin to derive our method.

Overview. The Dincta algorithm inputs a PCA embedding (*Z*) of cells, along with their batch assignments (Φ_*B*_), the all, partial or none cell type assignments (Φ_*C*_), the unknown cell type indicator (*u*) of cells and the scaling factor *α* for the cells with unknown cell type. This algorithm was summarized as Algorithm 1 below. The outer loop updates the unknown cell type indicator (Algorithm 6) until it converges by checking the fitness between the cell type assignments and cluster assignments; the inner loop iterates between two complementary stages: maximum diversity clustering and inferring (Algorithm 2), and a mixture model based linear batch correction (Algorithm 5). The clustering and inferring step use the batch corrected embedding 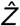 to compute a soft assignment of cells to the clusters, encoded in the matrix *R*, and then use the cluster assignment matrix *R* to update the cell type assignment matrix Φ_*C*_. The correction step uses these soft clusters to compute a new corrected embedding from the original one. Efficient implementations of Dincta, including the clustering, inferring and correction subroutines, are available as part of the R package at https://github.com/songtingstone/dincta.

### Algorithm 1 Dincta

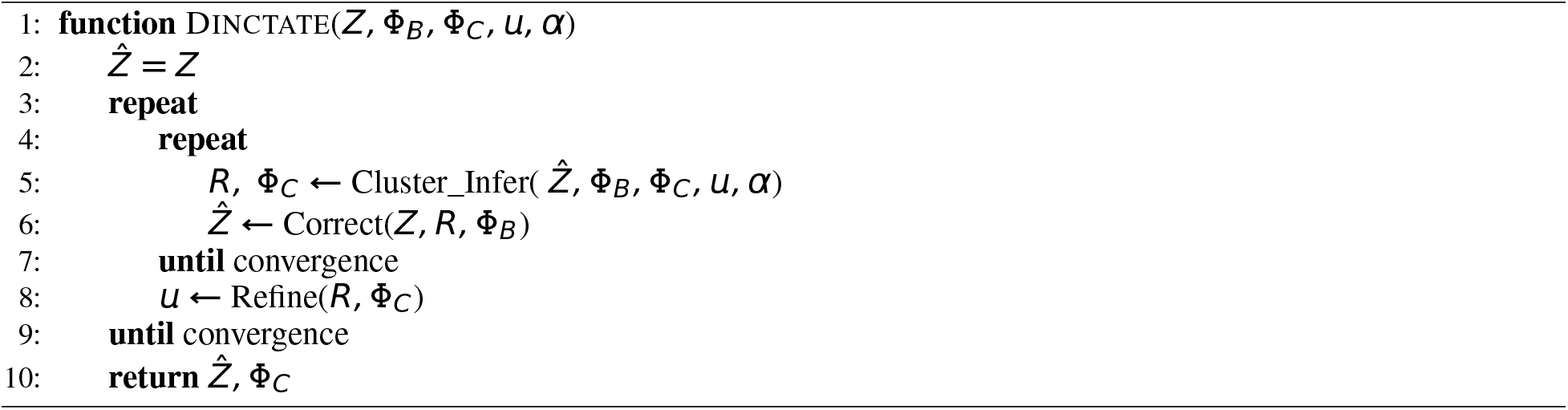

### Algorithm 2 Maximum Diversity Clustering And Inferring

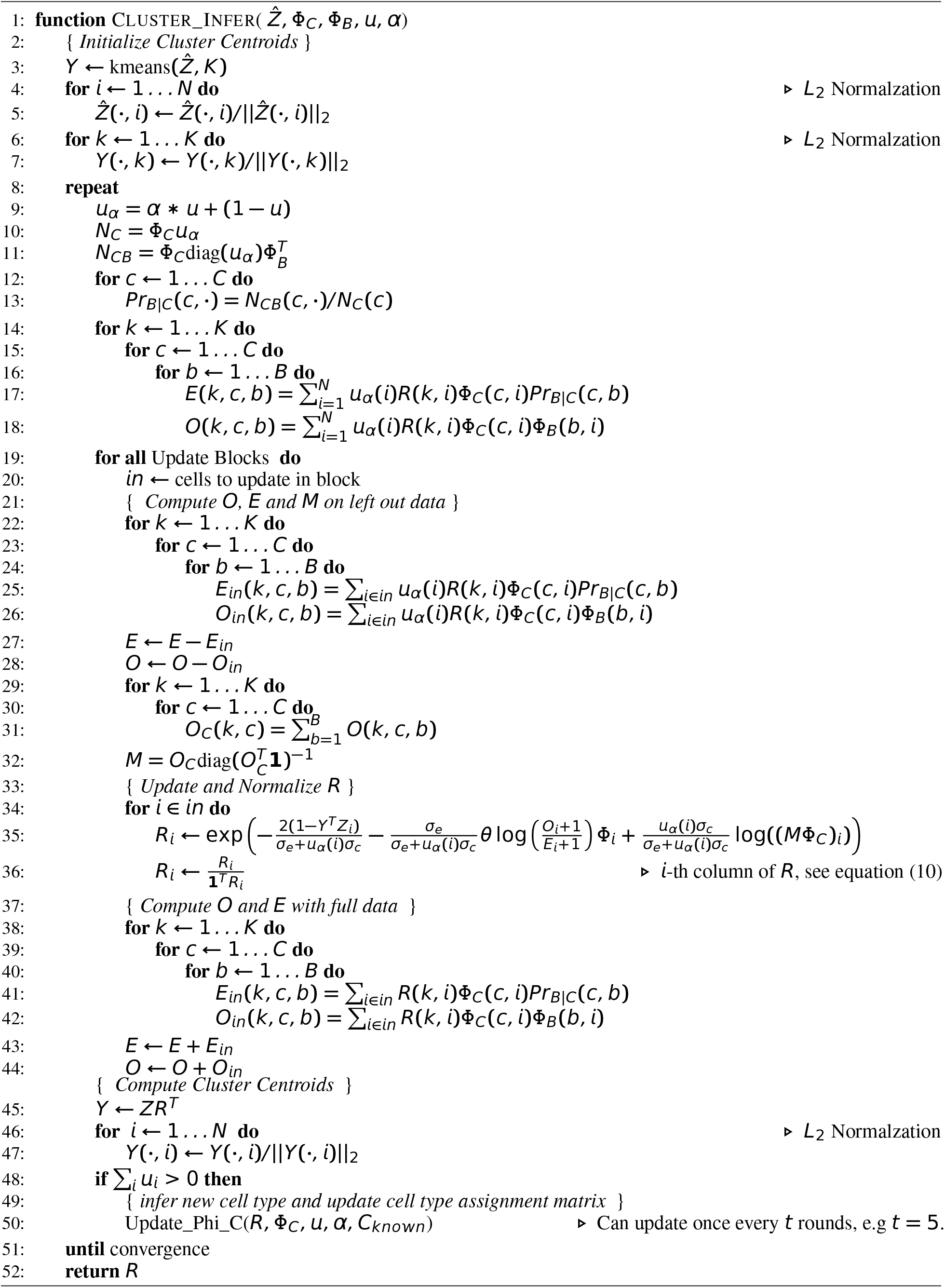

Glossary. For reference, we define all data structures used in all Dincta functions. For each one, we define its dimensions and possible values, as well as an intuitive description of what it means in context. The dimensions are stated in terms of *d*, the dimensionality of the embedding (for example, number of PCs); *B*, the number of batches; *C*, the number of cell types (containing the “unknown” cell type and new cell types); *N_b_*, the number of cells in batch *b*; and *K*, the number of clusters.

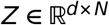 The input embedding, to be corrected in Dincta, This is often PCA embeddings of cells.

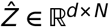 The integrated embedding, output by Dincta.

*R* ∈ [0, 1]^*K×N*^ The soft cluster assignment matrix of cells (columns) to clusters (rows).

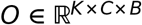 The observed co-occurrence array of cells in clusters, cell types and batches. For the cells with “unknown” cell type, we scale its contribution by a scaling factor *α*.

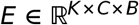 The expected co-occurrence array of cells in clusters, cell types and batches. For the cells with “unknown” cell type, we scale its contribution by a scaling factor *α*.

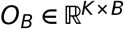 The observed co-occurrence matrix of cells in clusters and batches. For the cells with “unknown” cell type, we scale its contribution by a scaling factor *α*.

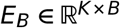 The expected co-occurrence matrix of cells in clusters and batches. For the cells with “unknown” cell type, we scale its contribution by a scaling factor *α*.

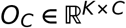 The observed co-occurrence matrix of cells in clusters and cell types. For the cells with “unknown” cell type, we scale its contribution by a scaling factor *α*.

*Y* ∈ [0, 1]^*d×K*^ Cluster centroid locations in the *k*-means clustering algorithm.

Φ_*B*_ ∈ [0, 1]^*B×N*^ The batch assignment membership of cells.

Φ_*C*_ ∈ [0, 1]^*C×N*^ The cell type assignment membership of cells. If there are cells with “unknown” cell type, then *C* = *C_Original_* + 1 + *C_new_* where *C_Original_* is the number of cell type known in the dataset. 1 accounts for the special “unknown” cell type, and *C_new_* is the number of new cell types in the data. For the initial input Φ_*C*_ of Dincta, *C_new_* = 0. And if the cell type of *i*th cell is unknown, then 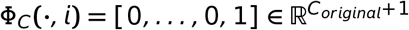, i.e the last row of Φ_*C*_ is for the “unknown” cell type. As the progress of the Dincta, it will infer the the number of new cell types *C_new_*, soft assign each cell with known or novel cell types, and also reduce the “unknown” cell type row Φ_*C*_(*C_original_* + 1, ·) to **0**. If the cell types of all cells are known initially, then *C = C_Original_* and there is no inference step.

*u* ∈ {0, 1}^*N*^ The indicator vector of “unknown” cell type. *u_α_* = 1 if the *i*-th cell’s cell type is unknown. *u_i_* = 0 if the *i*-th cell’s cell type is known. When we have the whole cell type information, then *u* = **0**.

*α* ∈ [0, 1] The scaling factor for the cells which have unknown cell type.

*u_α_* ∈ {*α*, 1}^*N*^ := *αu* + (1 − *u*) The scaling vector. *u_α_*(*i*) = 1 if the cell type of the *i*-th cell is known. *u_α_*(*i*) = *α* if the cell type of the *i*-th cell is unknown. It was used in the loss scaling and *E, O* and *M* computation, reduce the weight of information coming from the cells which have “unknown” cell type.

Φ ∈ [0, 1]^*C×B×N*^ The scaled cell type and batch assignment membership of cells, Φ(*c, b, i*) := Φ_*C*_(*C, i*)Φ_*B*_(*b, u*)*u_α_*(*i*).

*M* ∈ [0, 1]^*K×C*^ The condition probability *Pr*(*k|c*) matrix of cells in clusters (columns), cell types (rows).

*H* ∈ [0, 1]^*C×K*^ The condition probability *Pr*(*c|k*) matrix of cells in clusters (rows), cell types (columns).

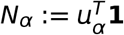 The effective count of cells.

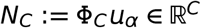 The effective count vector of cells in cell types.

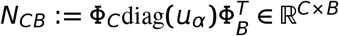 The effective observed co-occurrence matrix of cells in cell types and batches.

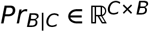 The probability matrix of the *Pr*(*b|c*).

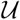 The index of cells with unknown cell type.

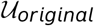 The index of cells with unknown cell type in the initial state.

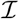 The index of cells with known cell type.

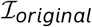 The index of cells with known cell type in the initial state.

Assumptions about the input data. In this article, we always use Dincta on a low-dimension embedding of the cells. To be clear about the properties of low-dimensional embedding space, we explicitly state three assumptions as described in Korsunsky et al. (2018):

1. Cells are embedded into a low-dimensional space as the result of PCA. The PCA embedding captures the variation of gene expression in a compact orthonormal space. For this reason, the default input to Dincta is a matrix of gene expression data, normalized for library size. We then perform PCA on this high-dimensional matrix and use the eigenvalue-scaled eigen vectors as the low-dimensional embedding input to Dincta.
2. Gene expression has been normalized for library size. In RNA-seq, each cell will be sequenced to a different depth, which results in different library sizes for each cell. It is best practice to account for this source of technical variation before performing PCA. In this article, we use the standard log counts-per10,000 (CP10K) transformation. As a result of the depth transformation, expression values are turned into relative frequencies inside each cell. Thus, it is impossible for every gene to be up-regulated in one group of cells.
3. The low-dimensional nearest-neighbor structure induced by Euclidean distance should be preserved with common similarity metrics such as cosine similarity and correlation. This can be easily checked by computing sparse nearest-neighbor graphs and comparing the adjacency matrices. Cells with less than 20% overlapping neighbors can be removed as outliers.

### Maximum Diversity Clustering and Inferring

We developed a clustering algorithm to maximize the diversity among batches within clusters and do not mix cells with different cell types. After we get the soft cluster assignment matrix, we use it to update cell type assignment matrix (inferring). We present this method as follows.

First, we review a previously published objective function for soft *k*-means clustering. Second, we add a diversity maximizing regularization term to this objective function, and derive this regularization term as the penalty on the statistical conditional dependence between two random variables: batch membership and cluster assignment, when given the cell type membership. Third, we add a penalty that constraint the the cluster membership in accordance with the cell type membership. We then derive and present pseudocode for an algorithm to optimize the objective function.

If there are cells with unknown cell type, we derive and present pseudocode to infer the known or novel cell type of cells with the cluster assignments.

#### Background: entropy regularization for soft k-means

The basic objective function for classical *k*-means clustering, in which each cell belongs to exactly one cluster, is defined by the distance form cells to their assigned centroids.

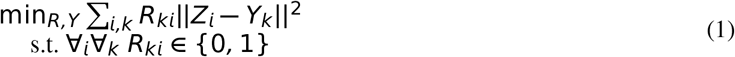

Above, *Z* is some feature space of the data and *Y* is centroids matrix. *R_ki_* denotes the membership of cell *i* in cluster *k* and take values in 0 and 1. To transform this into a soft clustering objective, we follow the derivation of Korsunsky et al. (2018) and add an entropy regularization term over *R*, weighted by a hyperparameter *σ_e_*. Now, *R_ki_* can take values between 0 and 1 with the constraint that *R_i_* must be a proper probability distribution with support [1, *K*].

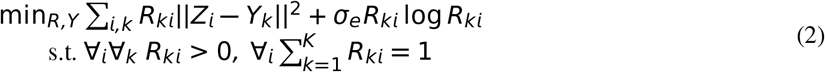

As *σ_e_* approaches 0, this penalty approaches hard clustering. As *σ_e_* approaches infinity, the entropy of *R* over-weights the data-centroid distances and each data point is assigned equality to all clusters.

In addition to soft assignment regularization, the function below penalizes cluster with low batch-diversity in each cell type. This penally, derived in the following section, depends on the cluster, cell type and batch identities 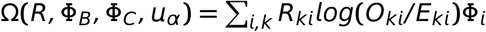.

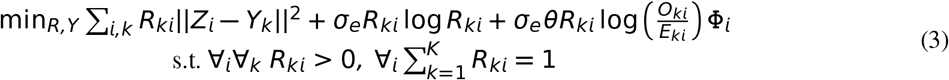

For each batch variable, we add a new parameter *θ*. The *θ* decides the degree of penalty for dependence between batch membership and cluster assignment in each cell type. When *θ* = 0, the problem reverts back to 2, with no penalty on the dependence. As *θ* increases, the objective function favors more independence between batch *b* and cluster assignment in each cell type.

#### Objective Function for Maximum Diversity Clustering

The full objective function for Dincta’s clustering builds on the previous section. In addition to soft assignment regularization and encourage batch-diversity in each type, we constrained the *R* satisfied the cell type probability relationship 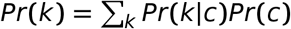 by penalizing the the Kullback Leibler divergence between *R* and *M*Φ_*C*_.

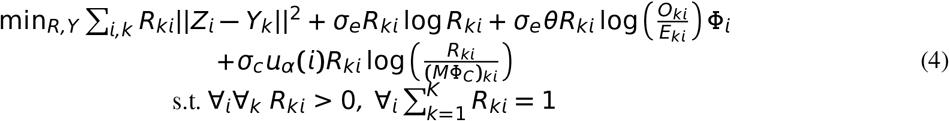

We add a new parameter *σ_c_*. Here the *σ_c_* decides the degree of penalty for the consistence between batch membership *R* and batch membership inferred from the probability relationship *M*Φ_*C*_. When *σ_c_* = 0, the problem reverts back to 3, with no penalty on the discrepancy between batch membership *R* and batch membership inferred from the probability relationship *M*Φ_*C*_. As *σ_c_* increases, the objective function favors the cluster assignments to be close to the cluster assignments inferred from the cell type assignments for each cell. As *σ_c_* approaches infinity, it will yield a hard constraint *R* = *M*Φ_*C*_.

For saving the computation power, we use the cosine distance instead of Euclidean distance, between *Z* and *Y*, which will constrain the clustering problem to the sphere clustering problem. Note that the squared Euclidean distance is equivalent to cosine distance when the vectors are *L*_2_ normalized. Therefore, assuming that all *Z_i_* and *Y_k_* have a unity *L*_2_ norm, the squared Euclidean distance above can be re-written as a dot product.

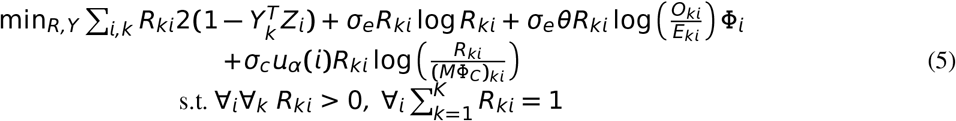

#### Cluster Diversity Score

Here, we discuss and derive the diversity penalty term Ω(·), defined in the previous section. For simplicity, we discuss diversity with respect to a single batch variable, as the multiple batch penalty terms are additive in the objective function. The goal of Ω(·) is to penalize the statistical dependence between batch identity and cluster assignment in each cell type. The observed co-occurrence counts (*O*) are compared to the counts expected under independence (*E*) for each cell type. We use the Kullback Leibler divergence (*D_KL_*), an information theoretic distance between two distributions to define the penalty. In this section, we define the *O* and *E* distributions, as we as the *D_KL_* penalty, in the context of the probabilistic cluster assignment matrix *R*.

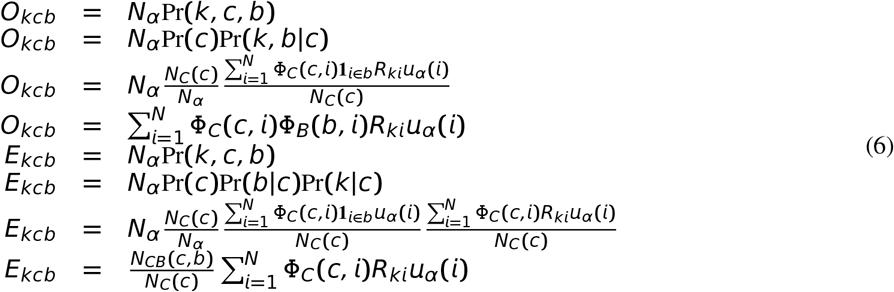

Next, we define the KL divergence in terms of *R*. Note that both *O* and *E* depend on *R*. However, in the derivation below, we expand one of the *O* terms. This serves a functional purpose in the optimization procedure, described later. Intuitively, in the update step of *R* for a single cell, we compute *O* and *E* on all the other cells. In this way, we decide how to assign the single cell to clusters given the current distribution of batches, clusters and cell types.

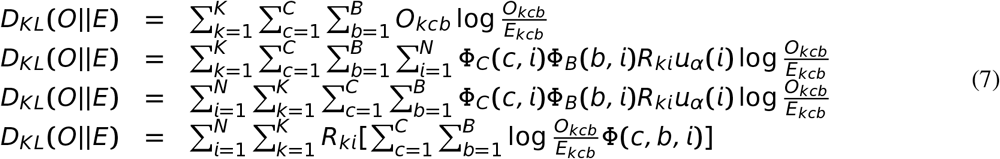

Define

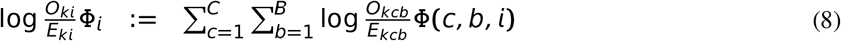

we can simplify the above equation to the following equation.

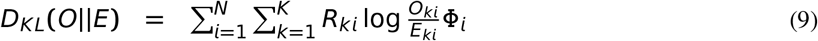

#### Optimization

Optimization of equation (5) admits an expectation-maximization framework, iterating between cluster assignment (*R*) and centroid (*Y*) estimation.

We solve for the optimal assignment *R*, for each cell *i*. First, we set up the Lagrangian with dual parameter λ and solve for the partial derivative with respect to each cluster *k*.

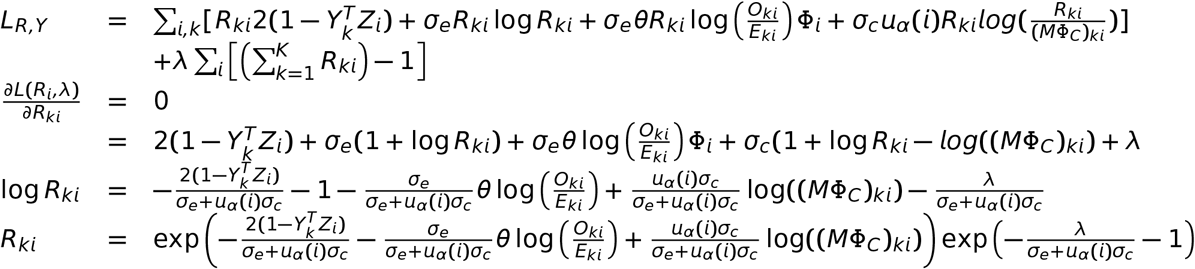

Next, we use the probability constraint 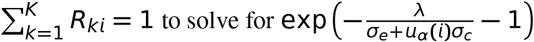

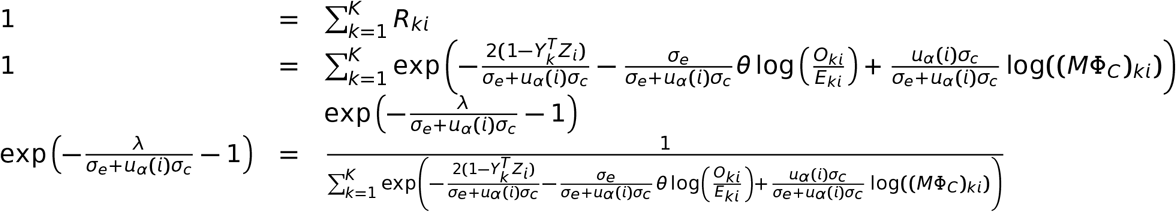

Finally, we substitute 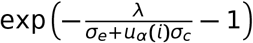 to remove the dependency of *R_ki_* on the dual parameter λ.

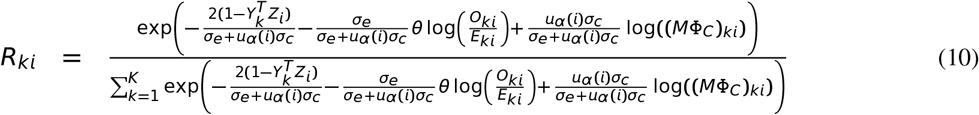

The denominator term above makes sure that *R_ki_* sums to one. In practice (Algorithm 2), we compute the numerator and divide by the sum.

Centroid estimation *Y*. Our clustering algorithm use cosine distance instead Euclidean distance. In the context of *k*-means clustering, this approach was pioneered by Dhillon and Modha (2001). We adopt their centroid estimation procedure for our algorithm. Instead of just computing the mean position of all cells that belong in cluster *k*, this approach then *L*_2_ normalizes each centroid vector to make it unit length.

The unnormalized expectation fo centroid position for cluster *k* would be 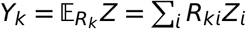. In vector form, for all centroids, this is *Y = R^T^Z*. The final position of the cluster centroids is given by this summation followed by *L*_2_ normalization of each centroid. This procedure is implemented in Algorithm 2 in the section {*Compute Cluster Centroids*}

Our algorithm can be run in two modes. In the first mode, we have the full cell type information of all cells. And we only need to find the cluster assignments *R* and correct the embedding *Z* by *R*. In the second mode, there are some or all cells with unknown cell type, and our algorithm can infer the known or novel cell type of cells with “unknown” cell type initially from cluster assignments *R*. In both kind of modes, the procedure of updating *R* is the same. Since at this time, we feed the Φ_*C*_ to the procedure as it as the true cell type assignments matrix. Now we give the maximum diversity clustering and inferring algorithm in Algorithm 2 and left the deduction of function update_Phi_C later.

#### Implementation details

The update steps of *R* and *Y* derived above form the core of maximum diversity clustering and inferring, outlined as Algorithm 1. This section explains the other implementation details of this pseudocode. Again, for simplicity, we discuss details related to diversity penalty terms *θ, Φ_B_, O* and *E* for each batch variable independently.

##### Block updates of *R*

Unlike in regular *k*-means, the optimization procedure above for *R* cannot be faithfully parallelized, as the values of *O, E* and *M* changed with *R*. The exact solution therefore depends an online procedure. For speed, we can coarse grain this procedure and update *R* in small blocks (for example, 5% of the data). Meanwhile, *O, E* and *M* are computed on the held out data. In practice, this approach succeeds in minimizing the objective for sufficiently small block size. In the algorithm, these blocks are included as the Update Blocks in the for loop.

##### Centroid initialization

We initialize cluster centroids by using regular *k*-means clustering, implemented in the base *R k*-means function. We use ten random restarts and keep the best one. We then *L*_2_ normalize the centroids to prepare them for spherical *k*-means clustering.

##### Regularization for smoother penalty

The diversity penalty term *O_kcb_/E_kcb_* can tend toward infinity if there are no cells from batch *b*, cell type *c* and cluster *k*. This extreme penalty can erroneously force cells into an inappropriate cluster. To protect against this, we add 1 to *O* and *E* to ensure that the fraction is stable (1 + *O_kcb_*)/(1 + *E_kcb_*). In this way, we can reduce the variance in the sample estimation of the counts matrix.

If some cell type has too small number of cells, then the (1 + *O_kcb_*)/(1 + *E_kcb_*) may fluctuate a lot. We can use the (1 + *O_C_*(*k.b*))/(1 + *E_C_*(*k.b*)) to replace it, where *O_C_*(*k.b*) := ∑_*c*_ *O_kcb_*), *E_C_*(*k.b*) := ∑_*c*_ *E_kcb_*). This can reduce the variance a lot. And it is implemented in the Dincta by set the the parameter “R.cross.entropy.type=‘b’”

##### *K*, the number of clusters

The number of clusters *K* use in Dincta soft clustering should be set to a value that reflects the size and complexity of the dataset. Too few clusters will not capture the number of biologically distinct cell types and states. Too many clusters will give too much weight to batch-specific outliers and prevent effective integration. As a heuristic, we assume that the dataset have at most 100 distinct cell types and that each cluster should have at least 30 cells. We set the default number of clusters, *K*, to lie between these two extremes for *N* cells.

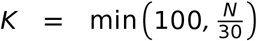

### Cell Type Inference

#### Cell type assignment and cluster assignment relation

Here, we discuss the probability relationship between cell type assignment and cluster assignment. And then we use their relations to infer the cell type of the cells. Let Pr(*k, c*) be the probability that cell type *c* and cluster *k* occur together, Pr(*k*) be the probability that cluster *k* occurs, and Pr(*c*) be the probability that cell type *c* occurs. Then we have the follow probability relationships

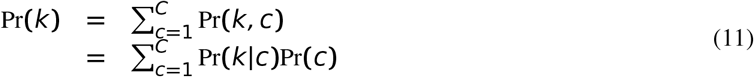

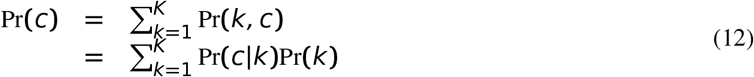

We can apply the above probability relation equation to each cell to get the probability relationship for each cell, which reveals how to transform between the cluster assignments and cell type assignments.

Before doing that, we prepare some necessary notations. Note that *R_ki_* is the probability that cell *i* assigned to the cluster *k*, Φ_*B*_(*b, i*) is the probability that cell *i* belong to batch *b*, and Φ_*C*_(*c, i*) is the probability that cell *i* assigned to the cell type *c*. With the three quantities, we can estimate the the transform probability matrix element Pr(*c|k*) by *H_ck_*.

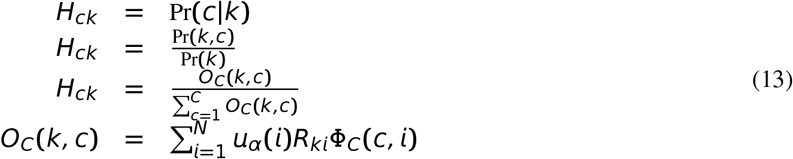

Similarly, we can estimate the the transform probability matrix element Pr(*k|c*) by *M_kc_*.

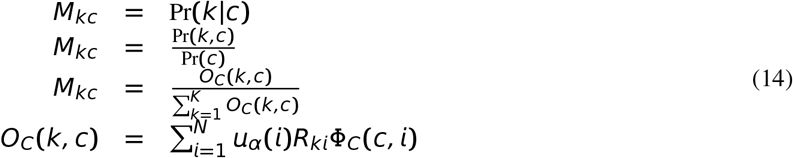

With these preparations, we apply the probability relation equation (11) to each cell. We get the following equation.

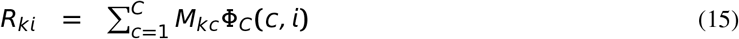

Similarly, we apply the probability relation equation (12) to each cell. We get the following equation.

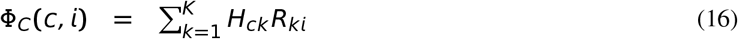

The above equations can be represented in the matrix form. Note that 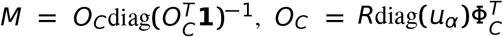, we can represent *M* by *R* and Φ_*C*_ with the following equation.

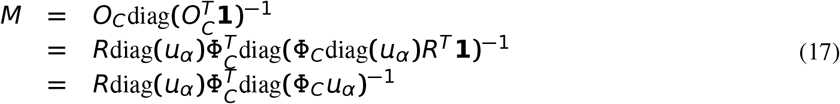

where the third equality comes from the *R* is a probability matrix which means the sum of each column of *R* is 1.

Similarly, note that 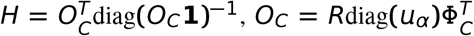, we can represent *H* by *R* and Φ_*C*_ with the following equation.

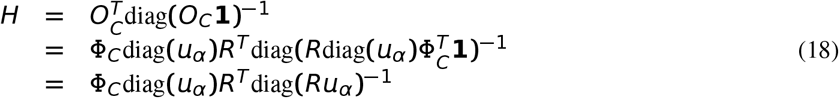

where the third equality comes from the Φ_*C*_ is a probability matrix which means the sum of each column of Φ_*C*_ is 1.

Combining the equation (15) and (17), we get the following equation.

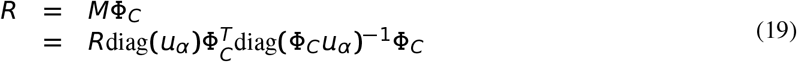

Similarly, combining the equation (16) and (18), we get the following equation.

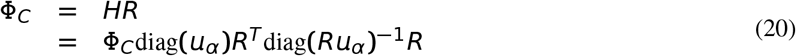

From equation (19) and (16), we see that *R* and Φ_*C*_ have very close relationship. To make the relation more clearly, we multiply diag^1/2^(*u_α_*) in the left and right side of equation (19) and (16). This give the following equations.

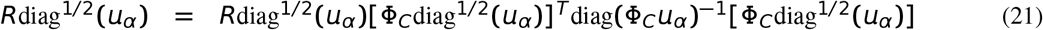

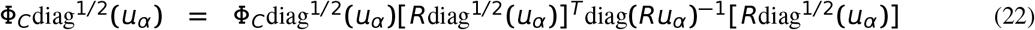

We list some of the property about *R*diag^1/2^ (*u_α_*), Φ_*C*_diag^1/2^ (*u_α_*) and their invari ant projection matrix *T_C_* := [Φ_*C*_diag^1/2^ (*u_α_*)]^*T*^diag(Φ_*C*_*u_α_*)^−1^ [Φ_*C*_diag^1/2^ (*u_α_*)] and *T_R_* := [*R*diag^1/2^ (*u_α_*)]^*T*^ diag(*Ru_α_*)^−1^ [*R*diag^1/2^ (*u_α_*)].

1. The invariant projection matrix *T_C_* of *R*diag^1/2^(*u_α_*) is symmetric semi-positive definite matrix. Its largest eigenvalue is 1 of algebraic multiplicity at least of rank of *R*diag^1/2^.
2. Each row of *R*diag^1/2^ is the right eigenvector of the matrix *T_C_* about the eigenvalue 1. 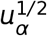 is the left eigenvector of *T_C_* about about the eigenvalue 1.
3. The invariant projection matrix *T_R_* of Φ_*C*_diag^1/2^(*u_α_*) is symmetric semi-positive definite matrix. Its largest eigenvalue is 1 of algebraic multiplicity at least of rank of Φ_*C*_diag^1/2^ (*u_α_*).
4. Each row of Φ_*C*_diag^1/2^ (*u_α_*) is the right eigenvector of the matrix *T_R_* about the eigenvalue 1. 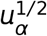 is the left eigenvector of *T_R_* about about the eigenvalue 1.

Let we look at the relationship between *R* and Φ_*C*_. *R* is the soft cluster assignment matrix, where the number of clusters is pre-specified and it usually larger than the number of the cell types. Φ_*C*_ is the cell type assignment matrix. And many clusters may be mapped to the same cell type. *R* is fine grained version of Φ_*C*_, may be viewed as the subtypes assignment matrix. So we can combine the information from *R* to infer the cell type of cells with unknown cell type initially by the formula Φ_*C*_(·, *i*) = *M^T^R*(*·, i*) if *M* is accurate. If we find the correct cluster assignment *R*, we also get the correct cell type assignment Φ_*C*_.

In the practice, the *M* is not an accurate estimation of Pr(*k|c*), since it depends on Φ_*C*_ which is not accurate. However, note that if we know the cell type assignments 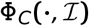 for the cells with known cell type initially and we have the correct cluster assignment matrix *R*, we can infer the cell type assignments 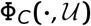 for the cells with unknown cell type by solving the linear equation (22). In details, we fetch out the columns indexed by 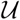 in the left and right side of (22), we get the following equation.

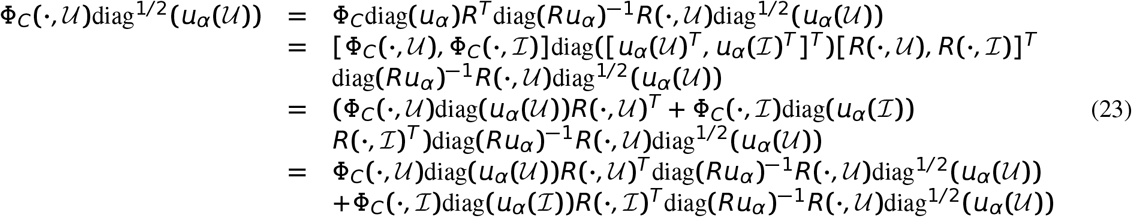

From above equation, we see that cell type assignment matrix 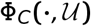 for the cells with unknown cell type satisfies the following linear system

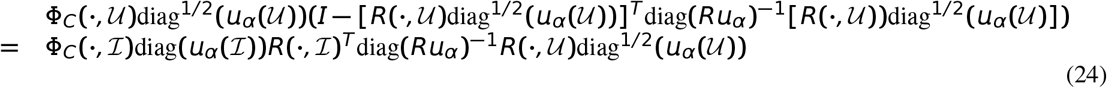

The matrix 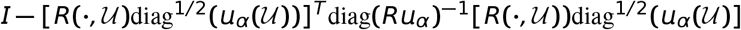 is symmetric semi-positive definite, and 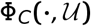 should be a probability matrix, i.e. 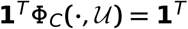. In generally, the solution is given by the following constrained least square problem.

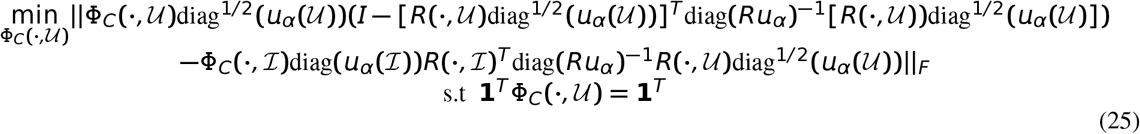

where 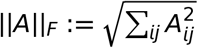 is the Frobenius norm of matrix *A*.

Note that the above problem is not easy to solve. Luckily, as in the following theorem, if *R* satisfies a mild condition, the least square problem without constraint has an explicit unique solution, and we can normalize the solution on each column to make it to the the probability matrix as an approximate solution of the constrained least square problem. We summarize it in the following theorem, and proof is in the supplementary.

##### Theorem 1.

*If* 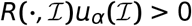 *elementwisely, then* 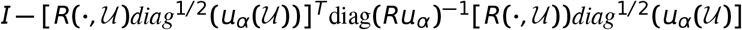 *is symmetric positive definite hence invertible and the least square problem*

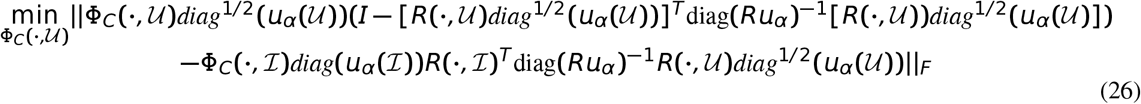

*has an unique solution*

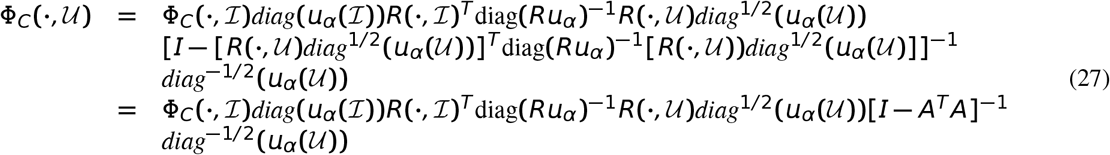

*and can be computed efficiently in the following representation*

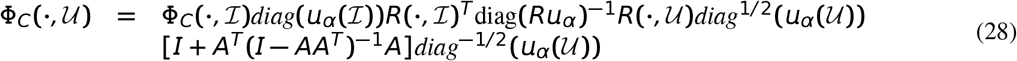

*where* 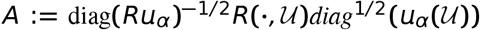. *And the solution of problem (26) can be approximated by normalize each column of* 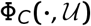 *to have mass* 1 *in equation (28). i.e*.

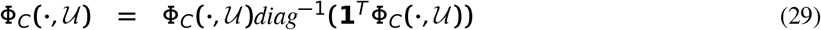

*In general, the matrix AA^T^ is semi-positive and the its largest eigenvalue λ_max_*(*AA^T^*) *is bound by max_k_β_k_, where* 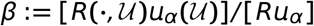 *and the operator / is element-wise division*.

If we have enough cells whose cell types are known, the cell types of all cells include in the cells with known cell type, and *R* finally converged to the correct cluster assignments, we can faithfully infer the unknown cell type’s cell type from the known cell type assignments matrix 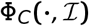 through solving the problem (26).

On the other side, if a new cell type in the data set occurs but we do not know initially, then the *R_ki_* will catch these new cell type with some new cluster *k*, and 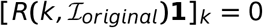. Then the assumption of Theorem 1 will be violated.

In this time, we can infer the new cell type assignment from 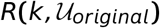 for the core cells of the new cell types. We can gain the intuition from a simple example. If there are two cell types, and cells with the two cell types are separate very well. In the first cell type *c* = 1, there are 2 clusters *k* = 1, 2, in the second cell type *c* = 2, there are 3 clusters *k* = 3, 4, 5. And suppose that the first cell type has *N*_1_ cells, and there are *N*_2_ cells are the second cell type. Then in the ideal case, *R* should be in the following form

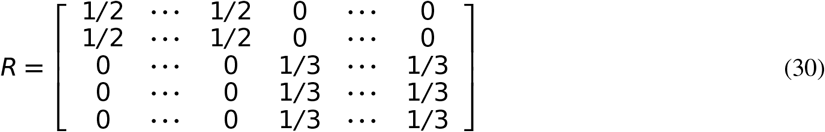

Now suppose that we don’t know the cell types information, i.e. 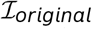 is empty, can we infer the number of cell types from the *R* matrix? the answer is yes! The number of cell types is the rank of matrix *R*, here is 2. We can get Φ_*C*_ by combining the rows of *R*, i.e., 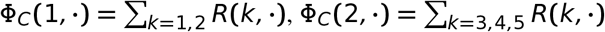. Indeed, this gives us an unsupervised method to classify cells with the help of the soft kmeans, which is implemented in DinctaClassfication function of Dincta package.

We generalize the above results in the following theorem as the foundation of Dincta to infer new cell types.

##### Theorem 2.

*Suppose that there are N cells belong to C cell types, and each cell type c has N_c_ cells. Moreover, each cell type separates very well, which means that* Φ_*C*_ ∈ {0, 1}^*C×N*^. *Suppose that Dincta algorithm output an ideal matrix R such that each cluster k only belong to one cell types c. Denote* 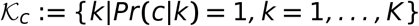, *and* 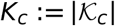. *And suppose that we can reorder the columns of R such that R has the following form*

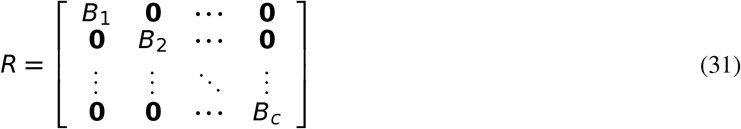

*where* 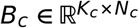

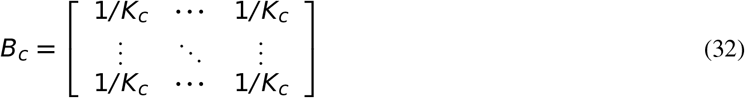

*and the corresponding* Φ_*C*_ *has the following form*

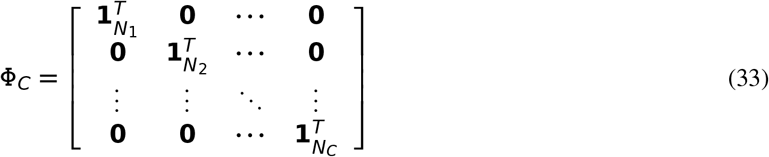

*where* **1**_*N_c_*_ *is the column vector whose element is* 1 *and its length is N_c_. Let A* : = *diag*^−1/2^ {*R***1**}*RR^T^diag*^−1/2^ { *R***1**}.

*Then A has the following form*

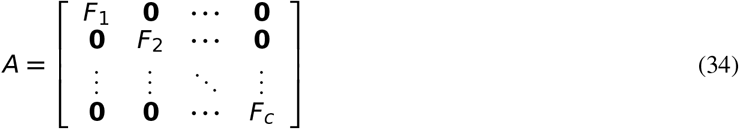

*where* 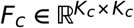 is

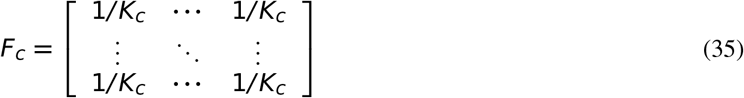

*The rank of* Φ_*C*_, *R, A are the same, i.e. rank*(Φ_*C*_) = *rank*(*R*) = *rank*(*A*) = *C. A is a symmetric semi-positive definite real matrix with eigenvalue* 1 *of algebraic multiplicity c and eigenvalue* 0 *of algebraic multiplicity K − C. Formally, the eigendecomposition of A has a special form* 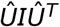, *where I* ∈ {0, 1}^*C×C*^ *is the identity matrix and* 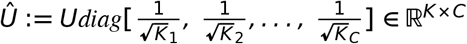 *is the columns orthogonal matrix whose columns are eigenvectors of A. And U is represented by*

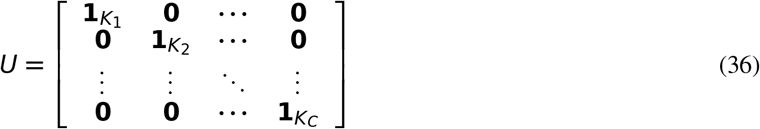

*Moreover*, Φ_*C*_ = *U^T^R, i.e. U is also the gather matrix that combines the rows of R to form the cell type assignment matrix* Φ_*C*_.

The proof of this theorem is simple and left to the reader.

In the above theorem, it gives us the intuition that the number of cell types is the algebraic multiplicity of eigenvalue 1 of the symmetric semi-positive definite real matrix *A*. And if one can find the special orthogonal eigenvectors matrix *U* of *A*, we can recover the cluster assignment matrix *R* to the cell type assignment matrix Φ_*C*_. As the assumption in the theorem, this phenomenon only occurs when that the cell type in the dataset is separate very well. In the reality, the dataset may be noisy, and also there are intermediate cell states which restrict the cluster assignment matrix do not satisfy the well separation assumption. Although there are intermediate states and cells in the boundary between two cell types, we can always find the core cells in the center of each cell type, and the core cells of each cell type satisfy the well separation assumption. We can select out those cells out and use the cluster assignments of the core cells to form the matrix *A*. Then *A* can determine the number of cell types and the gather matrix *U*. And the cell type assignments for the core cells can be obtained from *U* and *R*. Once we know the cell type assignments for the core cells with the new cell types, we regard these cell type assignments as known information. Combining with the already known cell type information, we can use the formula (28) and (29) to infer the cell types of the other cells on the boundary of cell types or the intermediate state cells.

In order to make the ideas to become computable, we must find an algorithm to compute the gather matrix *U*. Below, we elucidate the intuition to make this happen. We first use the eigenvalue algorithm to find the eigenvalues of *A* and determine the *C* as the rank of *A*. Note that the number of 1s in *U* is *K*, so we can compute the *U* by find a path that join (add) the columns of the identity matrix 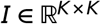 sequentially to get the *U*. To do this, we need fan objective to determine which two columns to join in each time.

We now first give our objective function, and then give a simple example to illustrate the key ideas.

Let the *U_K_ = I* be the start point of *U*, and 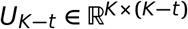 is the result after *t* times sequentially combining the columns of *U_K_*. And after *t = K − C* combination, we get that the gather matrix *U_C_ = U*. Suppose that we are in the step *t* ≥ 1, and currently we have the 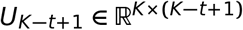, we want to find the index which two columns of *U*_*K−t*+1_ should be combined to get the 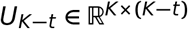. The key idea is that we find which is the smallest element of the off-diagonal elements of the objective matrix *D*_*K−t*+1_ := [*AU*_*K−t*+1_ − *U*_*K−t*+1_]^*T*^[*AU*_*K−t*+1_ − *U*_*K−t*+1_]. If there are multiple smallest elements, we select the element whose index is (*i, j*) such that *D*_*K*−1+1_ (*i, i*) + *D*_*K*−1+1_ (*j, j*) is the maximum among all these multiple smallest elements. And suppose that is element index is (*c*_1_, *c*_2_), then we should combine the *c*_1_, *c*_2_ of *U*_*K−t*+1_ by adding to get the *U_k−t_*.

To get some insights, let we think a simple case. Suppose that there is only one cell type, and this cell type is represented by *K* clusters. Now 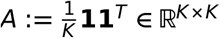, and we have 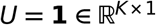. One can easily imagine that we can get *U* by sequentially combining the columns of initially identity matrix 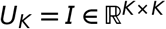 to get the final *U* := **1**. At step 1, *D_K_* = (*AI − I*)^*T*^ (*AI − I*) = (*AI − I*)^*T*^(*A − I*), one can easily verify that

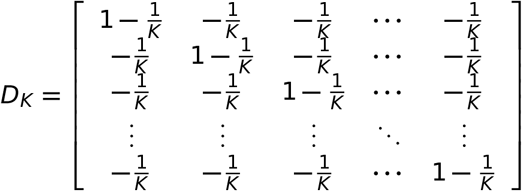

Note that each row sum, column sum and the sum of all elements of *D_K_* is zero, i.e. 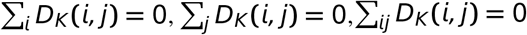. On this time we can random select two columns of *U_K_* to combine. For simplicity, we select the first two columns of *U_K_* to combine, this will give us the following *U*_*K*−1_:

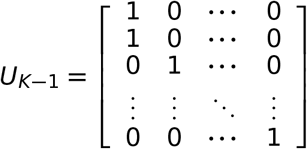

At step 2, *D*_*K*− 1_ = (*AU*_*K*−1_ − *U*_*K*−1_)^*T*^(*AU*_*K*−1_ − *U*_*K*−1_), one can easily verify that

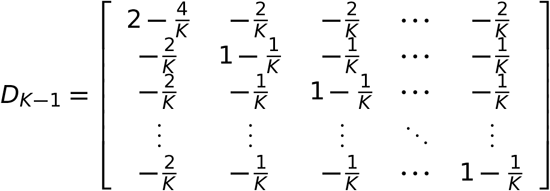

One can verify that the *D*_*K*−1_ can be obtained by combine the first two rows of *D_K_* by adding to get 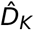 and then combine the fist two columns of 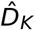 by adding to get *D*_*K*−1_. And in general, it is true, which can save our computation resources. Repeat above steps, we will finally get the *U* = **1** and *D*_1_ = **0**.

We summarize the above results in the following more general form of *A*, and the proof is simple but tedious and omits.

##### Theorem 3.

*Suppose that A has the following form*

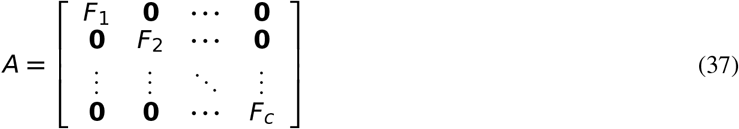

*where* 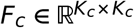 *is*

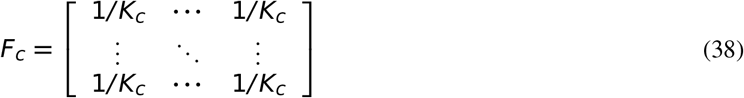

*Then A is a symmetric semi-positive definite real matrix with the eigendecomposition of* 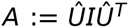, *where I* ∈ {0, 1}^*C×C*^ *is the identity matrix and* 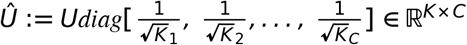 *is the columns orthogonal matrix whose columns are eigenvectors of A. And U is represented by*

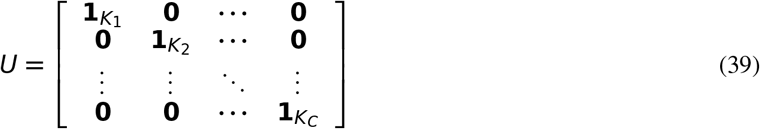

*We can get Û by the following algorithm. Let the U_K_ = I be the start point of U, and* 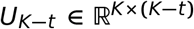 *is the result after t times sequentially combining the columns of U_K_. In the step t* ≥ 1, *we have* 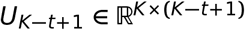. *Define the objective matrix D*_*K−t*+1_ := [*AU*_*K−t*+1_ − *U*_*K−t*+1_ ^*T*^[*AU*_*K−t*+1_ − *U*_*K−t*+1_]. *And let m be the minimum of the off-diagonal elements of D*_*K−t*+1_, *i.e. m* := min_*i≠j*_ *D*_*K*−1+1_ (*i, j*), *and let* 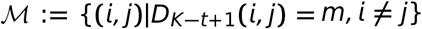 *be the index set achieved the minimum. We find the matrix index* (*c*_1_, *c*_2_) *such that* 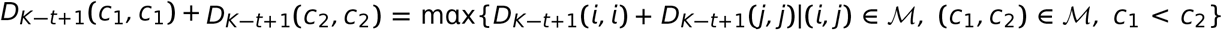. *U_k−t_ can be obtained by combining the c*_1_, *c*_2_ *columns of U*_*k−t*+1_ *by adding. And D_k−t_* := [*AU*_*K*−*t*_−*U_k−t_*]^*T*^ [*AU*_*K−t*_−*U_K−t_*] *can be obtained by combining c*_1_, *c*_2_ *columns of D*_*k−t*+1_ *to get* 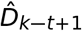, *and then combine the c*_1_, *c*_2_ *rows of* 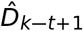 *by adding. And after t = K − C combinations, we get that the gather matrix U_C_ = U*.

We summarize above theorem in the Algorithm 3.

##### Algorithm 3 Find_Gather_U

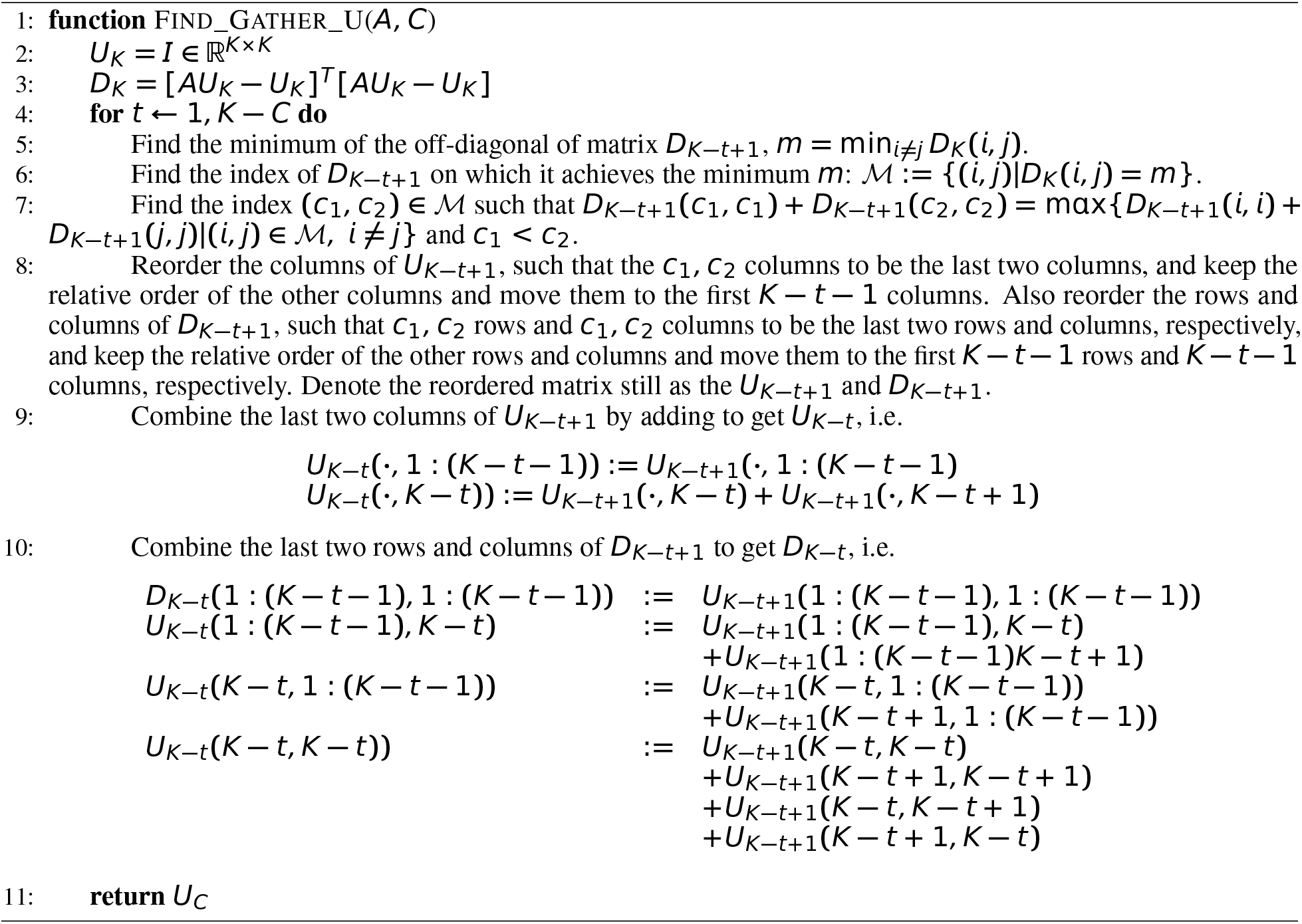

Now we can use this basic idea to infer the new cell types. Note that once there are new cell types in the cells with index 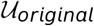, the soft kmeans clustering will catch the new cell types with some clusters. The indicator of this situations occuring is that 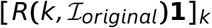 is near 0. In this case, we gather all new clusters 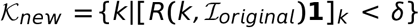, with a threshold *δ*. Usually *δ* = 6 is enough. We gather the index of the cells which belong to the new cell types to 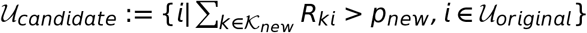 with the probability threshold *p_new_*, e.g. *p_new_* = 0.8.

In the progress of Dincta, the information in *R* is noisy, *R* is biased to the ideal cluster assignment matrix. To reduce the impact of the noise, we select cells from 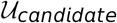 such that cells are in the center of the cluster it belongs to. We called these cells as the main cells, and they are gathered together to form the main set which is the index set of these main cells. We now explain the strategy used to gather the main set. The idea is very simple. We first estimate the effective number cells in cluster 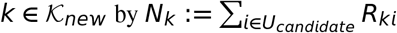. Then we set the select fraction *γ_s_* of *N_k_* for the each *k* set, respectively. For each 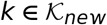, we sort the 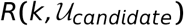 about the cell *i* with value *R_ki_* in descend order. And we select the top *γ_m_N_k_* cells ids to the main set, and finally join all ids to get the main set *U_m_*. As we shows later, we will fill the cell type assignment matrix 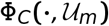.

To finish the inference of new cell type task, we must determine the number of new cell types and find the gather matrix *U* to transform the cluster assignment to the cell type assignment for the cells in the main set. And we can use the results of Theorem 3 to finish this.

Note that in the above simple example, we see that the number of the cell types is determined by the rank of the cluster assignment for those core cells which has a clear cell type, i.e. not on the boundary of two cell types. We now find a strategy to select the core cells from the whole data points, then we gather all these cells index into the sample set 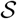.

To get the sample set, we use the strategy similar as we do to form the main set. We first estimate the effective number of cells in each cluster 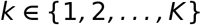 by 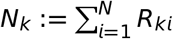. Then we set the the select fraction *γ_s_* of *N_k_* for the each *k* set, respectively. For each *k* ∈ {1, 2,…, *K*},we sort the *R*(*k, ·*) about the cell *i* in descend order. Then we select the top *γ_s_N_k_* cells ids to the sample set, and finally join all ids to get the sample set 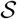.

After obtaining the sample set 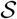, we compute the eigen decomposition of 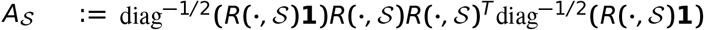. From the Theorem 3, the number of cell types is determined by algebraic multiplicity of eigenvalue 1 of 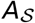. Note that there are noise in estimate *R*, we use the 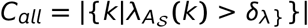 to determine of number of the cell types of the whole data, where *δ_λ_* ∈ (0, 1] is a threshold, e.g. *δ_λ_* = 0.9. Then we get the gather matrix 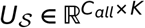 for 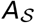 by the Algorithm 3.

To determine the number of new cell types *c_new_* and the corresponding gather matrix 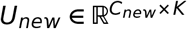 for the new cell types, we need to determine which columns of 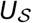 are the gather vectors for the new cell types. The indictor that one column of 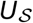 is the gather vector of new cell types is that 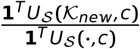 close to 1. And we denote the index of new cell type as 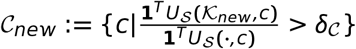 where 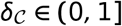 is threshold which determines the closeness to 1, usually 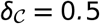 is enough. And the number of new cell types is given by 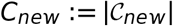. Finally, the gather matrix 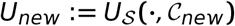, and 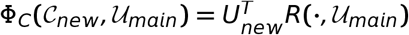 and 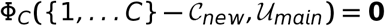.

Get all the efforts together, if the data contains new cell types, we can use the method described to find the main set of cells which belong to the new cell type. Then we get the cell assignment matrix for these main cells. After that, we combine the cell type information of the original cells 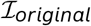 with the main cells with the new cell type 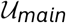 together, i.e. 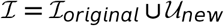. And the index of cells with unknown cell type reduced to 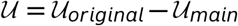. We can infer the cell type these cells by the equation (28) and (29).

We summarize the method which infers the cell type information matrix Φ*c* from the cluster assignment matrix *R* in the Algorithm 4.

##### Algorithm 4 Infer Cell Type Infomation

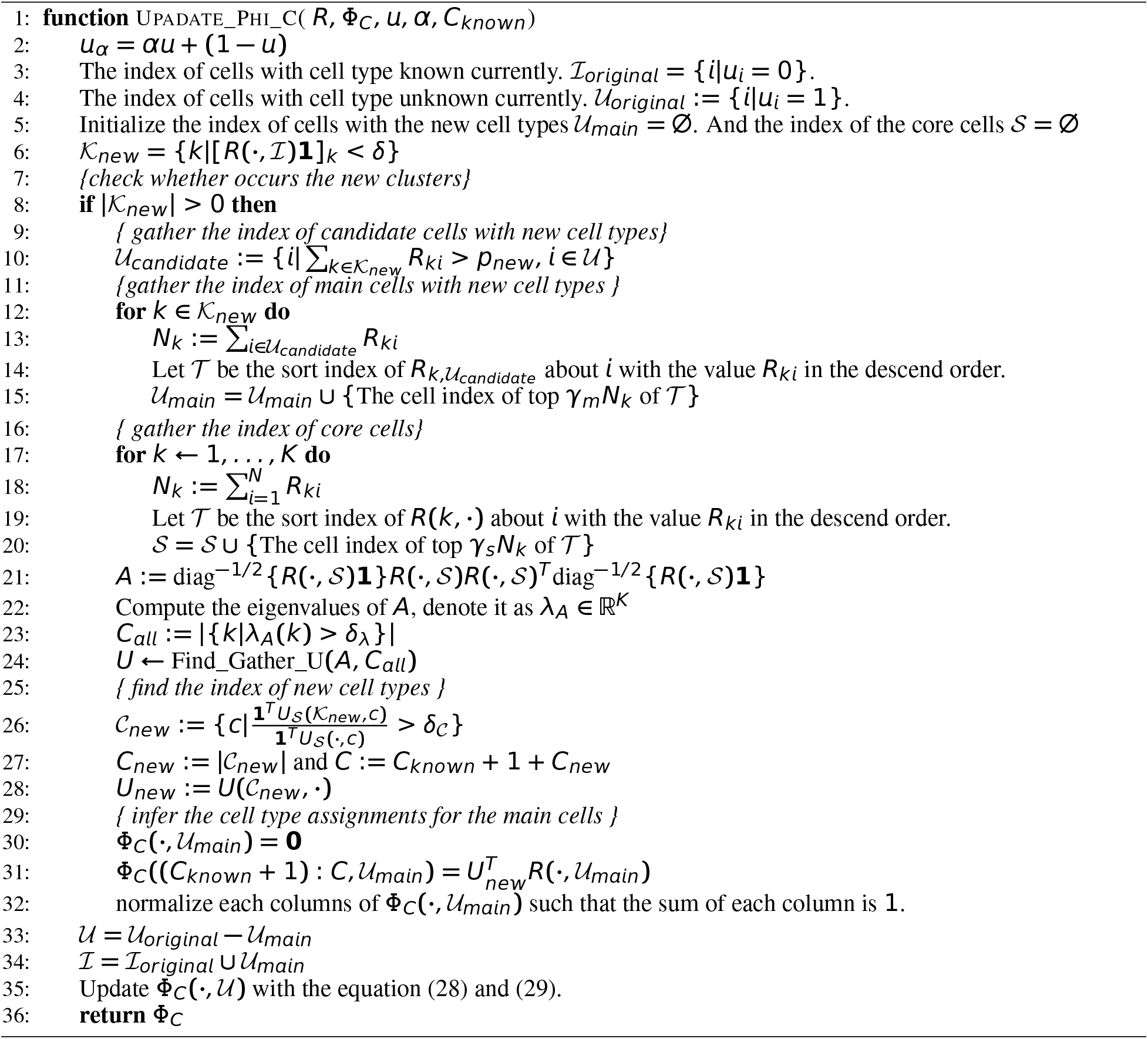

### Linear mixture model correction

We use the same correction model as in Korsunsky et al. (2018). To make the paper easy to read, we repeat the idea here.

In this section, we refer to all effects to be integrated out of the original embedding as batch effects. This dose not indicate that these effects are just technical. The terminology is only meant for convenient.

#### Mixture of experts model

Once we get the batch-diverse clusters assignments, we would like to remove batch-specific variation from each cluster. We achieve this with a variation on the original mixture of experts model by Jordan and Jacobs Jordan and Jacobs (1994). In the context of Dincta, each cell is probabilistically assigned to a small set of experts. This assignment was computed previously in the clustering and inferring step. Conditioned on a cluster/expert, mixture of experts assumes a linear relationship between the response and independent variables. Thus, we condition on cluster/expert *k* and define a Gaussian probability distribution for the response variables.

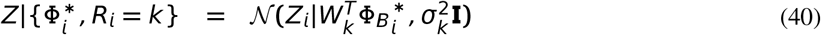

Here, the mean is 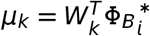, and the covariance is 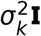 for simplicity. Note that the design matrix above 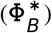 is not the same as the one used in the clustering step (Φ_*B*_). We augment the original design matrix Φ_*B*_ to include an intercept term: 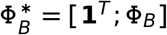. These intercept terms capture batch-independent (that is, cell type) variation in each cluster or expert. We can also achieve more complex behavior, such as reference mapping (Reference mapping) by modifying 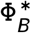.

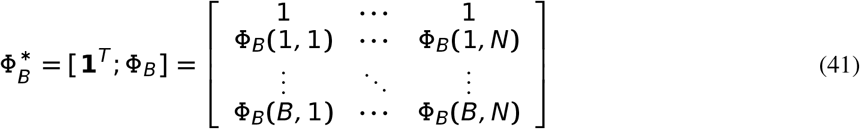

With this generative formulation, we can solve for the parameters (*W_k_*) of the linear model for each cluster or expert independently.

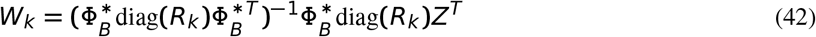

Above, diag(*R_k_*) is the diagonal matrix of cluster membership terms for cluster *k. Z* is the matrix of original PCA embeddings.

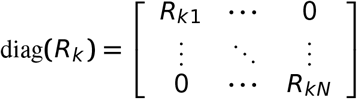

Each column in *W_k_* corresponds to a PC dimensioin, from *PC*_1_ to *PC_d_*. The first row of *W_k_* correpsonds to batch-independent intercept terms. The subsequent rows (1 to *B*) correspond to one-hot encoded batch assignments from the original design matrix Φ_*B*_, i.e. the batch effects vectors.

To get the cell-specific correction values, we take the expectation of *W_k_* with respect to the cluster assignment probability distribution *R*. In particular, each cell is modeled by a batch-independent intercept, represented in the first row of *W_k_*(*W*_*k*[0,]_) and its batch-dependent terms represented by the remaining rows *W*_*k*[1:*B*,]_.

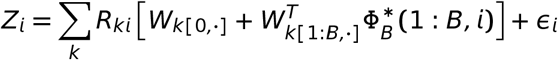

We split *W_k_* into two parts because we want to retain the intercept terms and remove the batch-dependent terms. To get the corrected embeddings 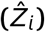, we subtract the batch-specific term 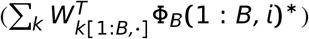 from the original embedding (*Z*).

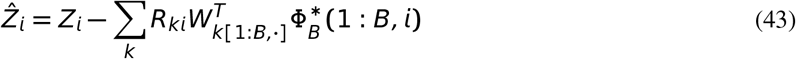

What remains is the cell-type specific intercept *W*_*k*[0,·]_ and the cell-specific residual *ϵ_i_*.

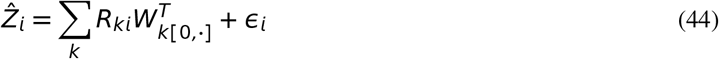

Unfortunately, for the design matrix 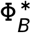, the formulation in equation (44) does not have a solution. This is because 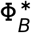 is not full rank and thus 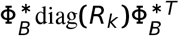 is not invertible. This singularity arise from the fact that the sum of a one-hot encoded categorical variable is equal to the intercept. To address this colinearity, we penalize nonintercept term in *W_k_* with an *L*_2_ norm, akin to ridge regression. This shrinks the *W_k_* terms to 0. Just as in ridge regression, instead of inverting 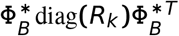, we invert 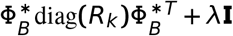, which is not singular, where *I* := diag([0, 1,…, 1]). The solution for *W_k_* now becomes

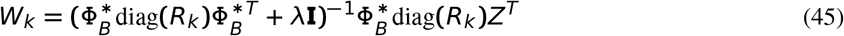

*λ* is the ridge penalty hyperparameter. Larger values of *λ* will shrink *W_k_* more toward 0. In practice, we set *λ* = 1. These correction steps are summarized in Algorithm 5.

##### Algorithm 5 Mixture of Experts Correct

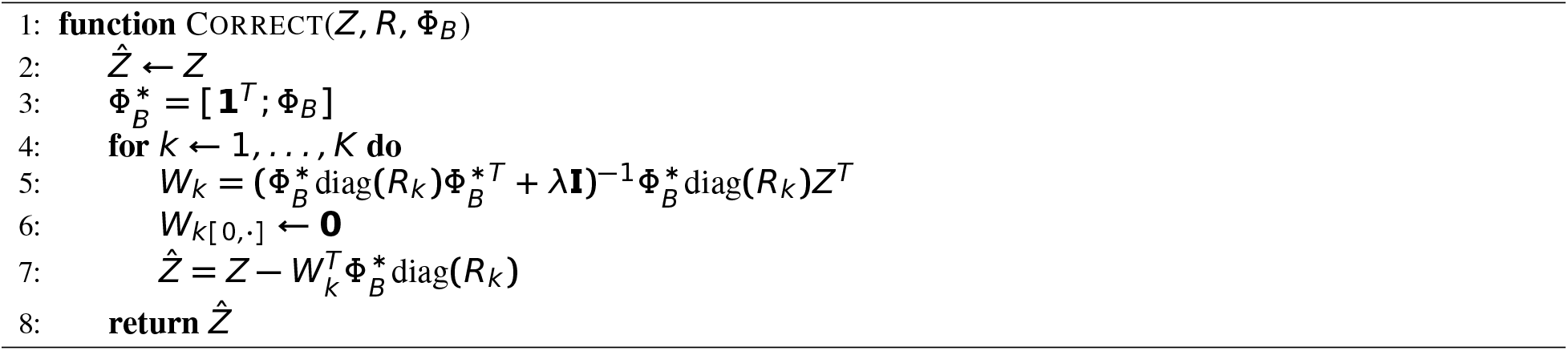

To understand that how this correction will move the embedding of the cells with the same cell type clustering together, we look at a very simple example. Suppose that there are only two batch, and each batch contains only one common cell type. And suppose that 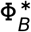 be in the following form:

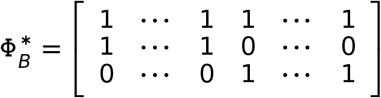

And 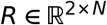 in the following ideal form.

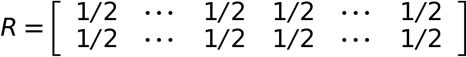

Now,

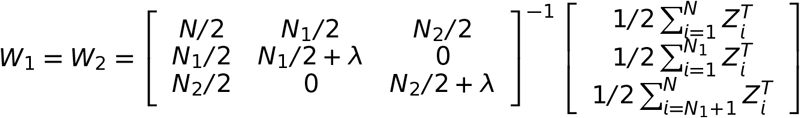

. Let *λ* = 1, with some mathematical calculations, we have

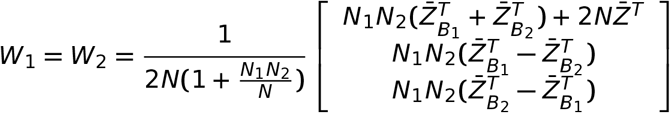

where 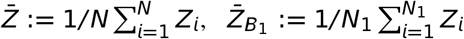 and 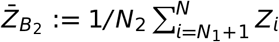. Suppose that *N*_1_ = *N*_2_ = *N*/2, then

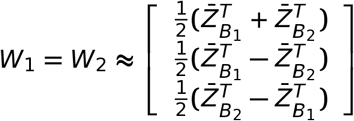

With the equation (43), the embedding of cells in the batch 1 *B*_1_ will be corrected by

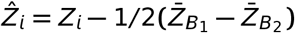

i.e. it will correct the embedding by move *Z_i_* toward the middle point 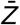 between the centroid 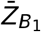 of batch 1 and centroid 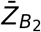 of batch 2. Similarly, the embedding of cells in the batch 2 *B*_2_ will be corrected by

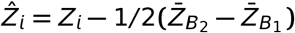

i.e. it will correct the embedding by move *Z_i_* toward the middle point 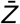 between the centroid 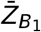 of batch 1 and centroid 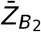 of batch 2. After correcting, the embedding of cells in the two batches will mix together, and move toward the center point 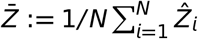.

#### Reference mapping

We can map query datasets onto a reference dataset, which is proposed in Korsunsky et al. (2018). We achieved this by modifying 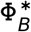 so that batch terms for reference cells do not get models or corrected. For every cell *i* in a reference dataset, set 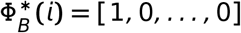. This makes Dincta explain cell *i* in terms of an intercept and nothing else. Since intercept terms do not get removed, cell *i* never changes its embedding (that is, 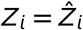).

#### Caveat

This section assumes the modeled data are orthogonal and each normally distributed. This is not true for the *L*_2_ normalized data used in the spherical clustering. Regression in this space requires the estimation and interpolation of rotation matrices, a difficult problem. We instead perform batch correction in the unnormalized space. The corrected data 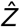 are the *L*_2_-normalized for the next iteration of clustering.

### Refine the cell type information

Once the inner loop converges in the Algorithm 1, we can update the unknown cell type information indicator vector *u* of cells. Here, we should find a criteria to identify which cell’s cell type is correct assigned. We do this by comparing KL divergence of Φ_*C*_ and 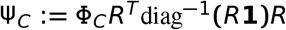 for each cell. In detail, the KL divergence distance of 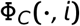 and Ψ_*C*_(·, *i*) is 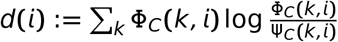. And we select the cells with the index 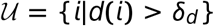 as the cells with unknown cell type, where *δ_d_* ∈ [0, ∞) is a threshold, usually *δ_d_* = 1*e* − 2 is enough. Then we update *u* with 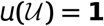 and 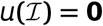 where 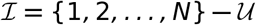. We summarize these steps in the Algorithm 6

#### Algorithm 6 Refine cell type assignment

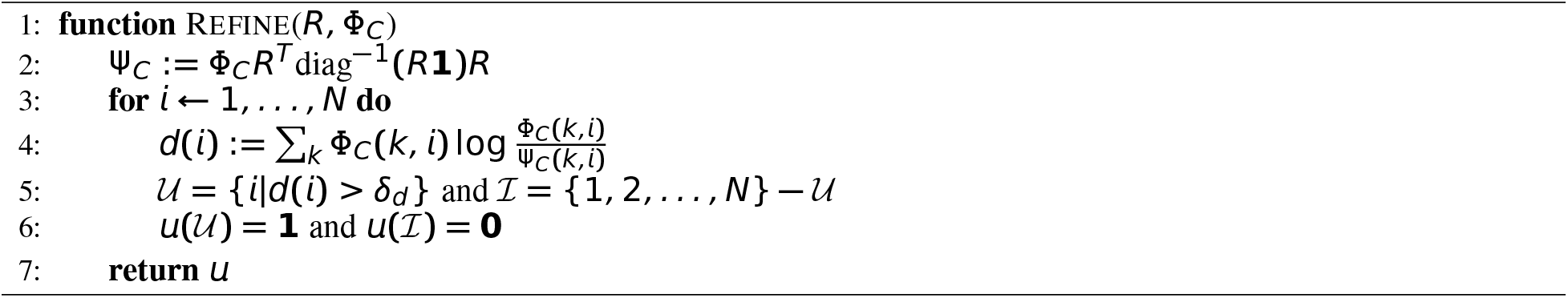

### Dincta Classification

Note that if there exists only one sample, and there is no need to do the batch correcting. We can shrink this model just by removing the batch variable and correcting step. This is implemented in Dincta by the function DinctaClassification. We omit the description of the algorithm and the numerical results for it to save space and make the main idea more clear.

### Infer Cell Type from the Cluster Assignment Matrix

To compare inferring accuracy, we can infer the number of cell types from directly from the cluster assignment matrix *R*. And this method also suit for any algorithm output a cluster assignment matrix.

The basic idea is that we define 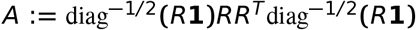, and the number of eigenvalues of *A* which are close to 1 as the number of cell types, i.e. 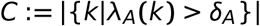. Moreover, if we have already know there are *C_known_* cell types, we set 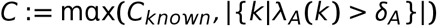. Then we use the find gather matrix algorithm 3 to find the gather matrix *U*, Finally, the cell type assignment matrix can be obtained by Φ_*C*_ : = *U^T^ R*. We summarize it in the following algorithm 7

### Performance and benchmarking

Assessing the degree of mixing during batch correction and dataset integration is an open problem. Several group have proposed methods to quantify the diversity of batches within local neighborhoods, defined by kNN graphs, of the embedded space. Butler et al. (2018) provide a statistical test to evaluate the degree of mixing, while Azizi et al. (2018) report the entropy of these distributions. Korsunsky et al. (2018) provide a interpretable metric LISI which is sensitive to the local diversity to quantify the diversity of mixing. In this paper, we use the LISI metric.

#### Match Score

We use the 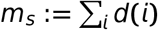 as the match score between the cell type assignment matrix and the cluster assignments matrix, where 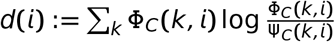 and 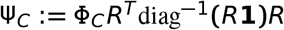.

#### Comparison Between Cell Type Assignment Matrix

Now, suppose that we have two cell type assignment matrix, one is the golden cell type assignment matrix Φ_*C*_ by the experts and the other is the inferred cell type assignment matrix Ψ_*C*_ such as from Dincta or output by the Algorithm 7, we propose the following algorithm 8 to compute the inferring accuracy of the Ψ_*C*_.

##### Algorithm 7 Infer Cell Type Information From Cluster Assignment Matrix

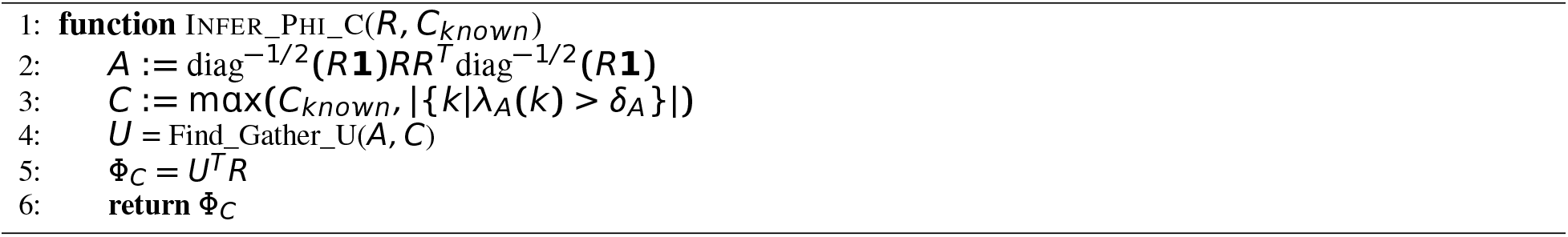

##### Algorithm 8 Inferring Accuracy

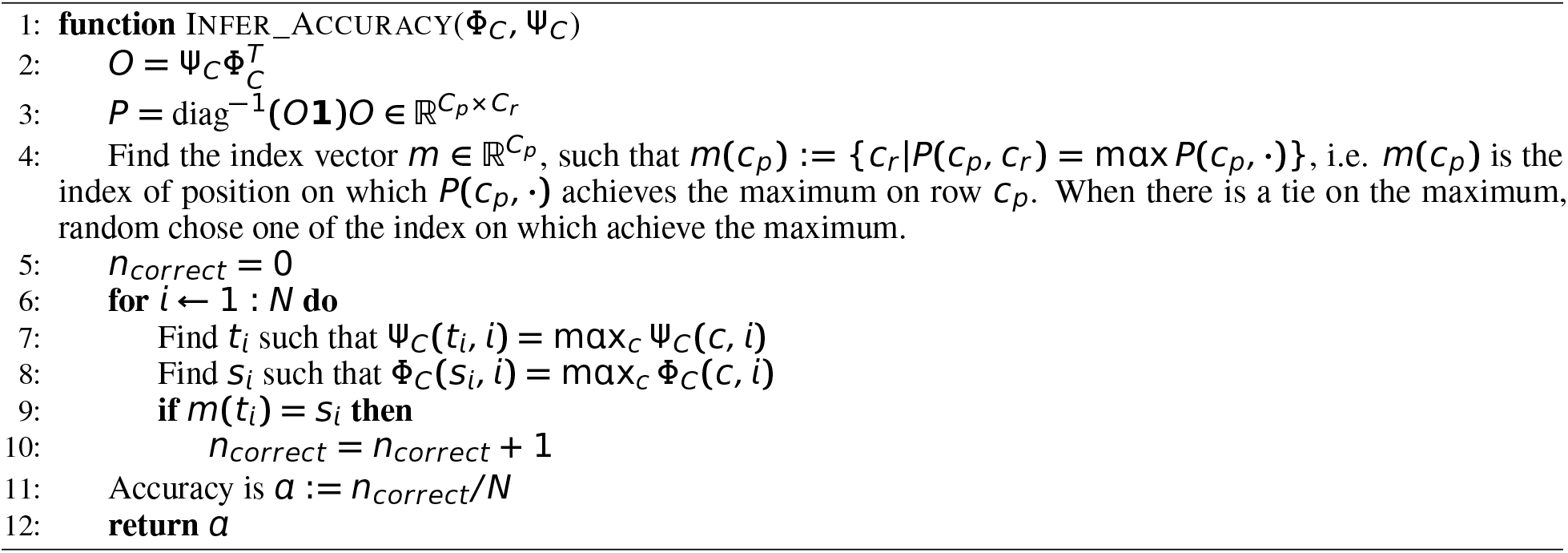

### Analysis details

#### Preprocessing scRNAseq data

Since we use the same data in Korsunsky et al. (2018), see the preprocessing steps in it.

#### Visualization

We used the UMAP algorithm to visualize cells in a two-dimensional space. For all analyses, UMAP was run with the following parameters: *k* = 30 nearest neighbors, correlation based distance and min_dist = 0.3.

#### Comparison to other algorithms

Note that he three datasets is analyzed in the Korsunsky et al. (2018) with the Harmony package and do the comprehensive comparison with other algorithm, such as MNN Correct (Lun et al., 2016), Seurat multiCCA (Butler et al., 2018), Scanorama (Hie et al., 2019), BBKNN (Polanski et al., 2019), it shows that Harmony do a better work on these datasets. For simplicity, we only compare the results of Dincta with the Harmony outputs.

#### Dincta parameters

By default, we set the following parameters for Dincta: *θ* = 2, *K* = 100, *τ* = 0, *σ_e_* = *σ_c_* = 0.1, block_size = 0.05, *ϵ*_cluster_ = 10^−5^, *ϵ*_Dincta_ = 10^−4^, max_iter_cluster_ = 200, max_iter_Dincta_ = 20, max_iter_refine_ = 3, *λ* = 1, *α* = 0.5, *δ* = 6, *P_new_* = 0.8, *γ_m_* = 0.6, *γ_s_* = 0.9, *δ_C_* = 0.5, *δ*_λ_ = 0.9, *δ_d_* = 1*e* – 2. For the pancreas analysis, we set donors to be the primary covariate (*θ* = 1) and technology secondary (*θ* = 2).

#### Harmony parameters

By default, we set the following parameters for Harmony: *θ* = 2, *K* = 100, *τ* = 0, *σ* = 0.1, block_size = 0.05, *ϵ*_cluster_ = 10^−5^, *ϵ*_Harmony_ = 10^−4^, max_iter_cluster_ = 200, max_iter_Harmony_ = 20, *λ* = 1. For the pancreas analysis, we set donors to be the primary covariate (*θ* = 1) and technology secondary (*θ* = 2).

## 5 Data availability

All data analyzed in this article are publicly available in the https://github.com/immunogenomics/harmony2019. And the original links of data can be found in Supplementary Table 8 in Korsunsky et al. (2018).

## 6 Code availability

Dincta are available as R package on https://github.com/songtingstone/dincta. Scripts to reproduce results of the primary analyses will be made available on https://github.com/songtingstone/dincta2020.

## 7 Supplementary

### Theorem 4.

*If* 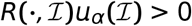 *elementwisely. then* 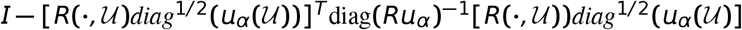 *is symmetric positive definite hence invertible and the least square problem*

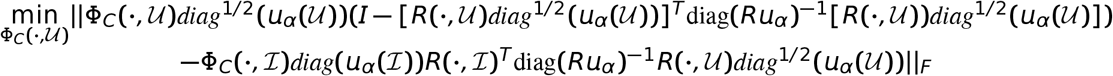

*has an unique solution*

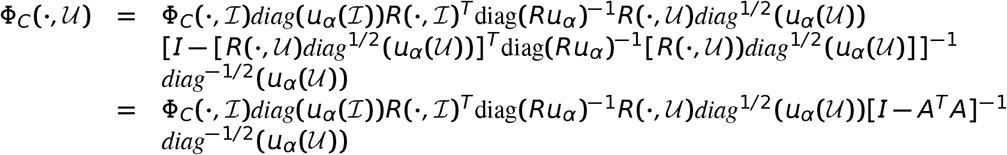

*and can be compute efficiently in the following representation*

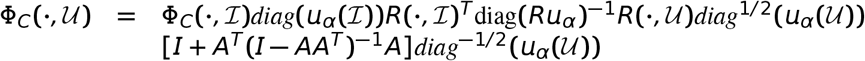

*where* 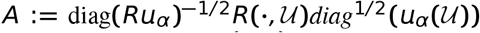. *And the solution of problem (26) can be approximated by normalize each column of* 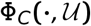 *in equation (28). i.e*.

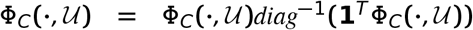

*In general. the matrix AA^T^ is semi-positive and the its largest eigenvalue λ_max_*(*AA^T^*) *is bound by max_k_β_k_. where* 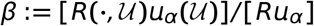 *and the operator / is element-wise division*.

*Proof*. We fist show that 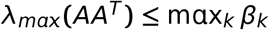, where *β*, where 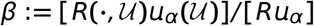. Let 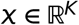 with the unit length, i.e. 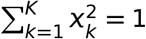. Note that 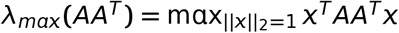.

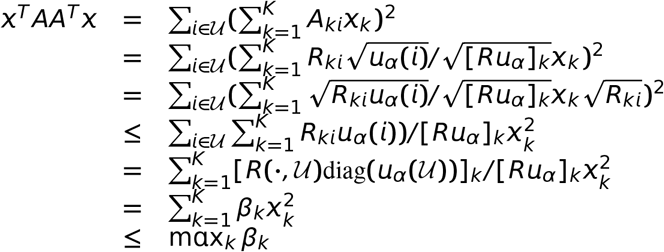

where the first inequality comes from the Cauchy inequality and 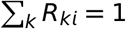.

Next, we show that if 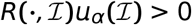, then 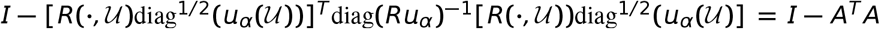 is positive hence invertible.

Note that if 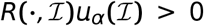, then 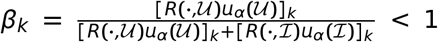. So we have 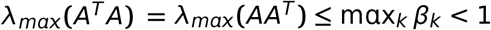. Then λ_*max*_(*I − A_T_A*) < 0, i.e. 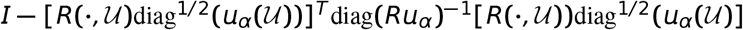 is positive hence invertible and the least square problem has the unique solution.

Finally, the equality (*I − AA^T^*)^−1^ = *I + A^T^*(*I − AA^T^*)^−lA^ can be proved by check that [*I − AA^T^*][*I − A^T^* (*I − AA^T^*)^−1^ *A*] − *I*, which is very easy to verify. And the proof is finished.

